# Disparate patterns of genetic divergence in three widespread corals across a pan-Pacific environmental gradient highlights species-specific adaptation trajectories

**DOI:** 10.1101/2022.10.13.512013

**Authors:** Benjamin C C Hume, Christian R Voolstra, Eric Armstrong, Guinther Mitushasi, Barbara Porro, Nicolas Oury, Sylvain Agostini, Emilie Boissin, Julie Poulain, Quentin Carradec, David A. Paz-García, Didier Zoccola, Hélène Magalon, Clémentine Moulin, Guillaume Bourdin, Guillaume Iwankow, Sarah Romac, Bernard Banaigs, Emmanuel Boss, Chris Bowler, Colomban de Vargas, Eric Douville, Michel Flores, Paola Furla, Pierre E Galand, Eric Gilson, Fabien Lombard, Stéphane Pesant, Stéphanie Reynaud, Matthew B. Sullivan, Shinichi Sunagawa, Olivier Thomas, Romain Troublé, Rebecca Vega Thurber, Patrick Wincker, Serge Planes, Denis Allemand, Didier Forcioli

**Author notes:** equally contributing (Benjamin C C Hume, Christian R Voolstra, Eric Armstrong, Didier Forcioli). **One sentence summary:** Coral evolutionary trajectories are species-specific. This is publication number 25 of the *Tara* Pacific Consortium.

## Abstract

Tropical coral reefs are among the worst affected ecosystems by climate change with predictions ranging between a 70-90% loss of reefs in the coming decades. Effective conservation strategies that maximize ecosystem resilience, and potential for recovery, must be informed by the accurate characterization of extant genetic diversity and population structure together with an understanding of the adaptive potential of keystone species. Here, we analyzed samples from the *Tara* Pacific Expedition (2016 to 2018) that completed an 18,000 km longitudinal transect of the Pacific Ocean sampling three widespread corals – *Pocillopora meandrina, Porites lobata*, and *Millepora* cf. *platyphylla* – across 33 sites from 11 islands. Using deep metagenomic sequencing of 269 colonies in conjunction with morphological analyses and climate variability data we can show that the sampled transect encompasses multiple morphologically cryptic species that exhibit disparate biogeographic patterns, and most importantly, distinct evolutionary patterns, despite exposure to identical environmental regimes. Our findings demonstrate on a basin-scale that evolutionary trajectories are species-specific and complex, and can only in part be predicted from the environment. This highlights that conservation strategies must integrate multi-species investigations to consider the distinct genomic footprints shaped by selection as well as the genetic potential for adaptive change.

## INTRODUCTION

Coral reef ecosystems harbor approximately one-third of the world’s multicellular marine biodiversity (Fisher et al. 2015) despite covering only ∼0.1% of the seafloor (Smith 1978). Critically however, these ecosystems are also some of the most sensitive to climate change (Gattuso et al. 2015) with global coral coverage estimated to have decreased by approximately 50% since the 1950s (Eddy et al. 2021). In line with even moderate projections of global warming, 70-90% of coral reefs may disappear in the coming decades (Allen et al. 2018), jeopardizing the biological diversity they support and the more than 500 million people who rely on the services they provide (Wilkinson 2008). To minimize further losses, and maximize their potential for recovery, the implementation of effective conservation strategies for these ecosystems, besides restoration/rehabilitation approaches, is imperative (Mcleod et al. 2019; Voolstra et al. 2021).

A critical component underlying the success of these conservation strategies is the preservation of the reef-building corals that form the ecological and physical foundations of reef ecosystems (Mcleod et al. 2019; Beger et al. 2014). Accurately characterizing the extant genetic diversity and genetic structure of these corals (Almany et al. 2009), as determined by their connectivity (Almany et al. 2009; Fontoura et al. 2022; Eric Pante et al. 2015), is an essential prerequisite for planning marine reserve networks that can replenish lost genetic diversity (Mcleod et al. 2019). Characterization of life history traits such as growth form and reproductive mode, and other biological attributes such as acclimation and adaptation potential are also essential, as different species may have significantly disparate responses to prevailing and historical environmental conditions (Selkoe et al. 2016; Buitrago-López 2021).

However, unrecognized interspecific (cryptic) diversity and high intraspecific morphological plasticity (e.g., in the genera *Acropora* and *Pocillopora*; see Flot, Couloux, and Tillier 2010; Richards, Berry, and van Oppen 2016; Pinzón et al. 2013) make characterizing genetic diversity through morphological characteristics alone problematic. The accurate characterization of diversity is improved through integration of morphological data with genome-wide sequencing strategies (e.g., RAD-seq, E. Pante et al. 2015; RNA-Seq, or whole genome/exome shotgun sequencing). Such approaches enable finescale resolution of genetic lineages, estimations of relatedness and divergence timings, and identification of genomic loci under selection (Carstens et al. 2013), all of which are of high value to conservation planning efforts. To best inform conservation efforts, robust genetic characterization of corals (i.e., from deep whole genome sequencing) across large geographic ranges are therefore required. Such campaigns are however difficult to achieve due to the logistical challenge of standardized sample collection and considerable financial requirements.

Across the Pacific Ocean it is largely unknown how coral diversity is structured, how the prevailing environment has shaped evolutionary history, and whether the consequential evolutionary trajectories are shared between corals. To begin to answer these questions, we analyzed samples from the *Tara* Pacific Expedition (Planes et al. 2019), that ran from 2016 to 2018 and completed an 18,000 km longitudinal transect of the Pacific Ocean, sampling three widespread corals – *Pocillopora meandrina, Porites lobata*, and *Millepora* cf. *platyphylla* – across 33 sites from 11 islands. We conducted ultra-deep metagenomic sequencing of 269 coral colonies in combination with morphological analyses and climate data to determine the standing genetic diversity and population structure of the coral species under study, identify several cryptic species, and reveal disparate patterns of environmentally linked genomic loci under selection. Our work demonstrates that different species are differentially shaped by the same prevailing environment and exhibit distinct genomic footprints. Such information needs to be taken into account to guide ongoing and future conservation efforts.

## RESULTS

Morphology-guided sampling of colonies resembling *P. meandrina, P. lobata*, and *Millepora* cf. *platyphylla* resulted in the collection of 106, 109, and 54 coral colonies across 11 islands, spanning 18,000 km of overwater distance (Figure 1; Supplementary Table 1). To resolve species identities, we designated the sampling regime as morphological-based primary species hypotheses (PSH) that we further developed into secondary species hypotheses (SSH) through an integrated taxonomic approach by analyzing genetic diversity and morphological parameters of the sampled colonies (Eric Pante et al. 2015). For each of the three coral genera, we used multiple genetic analyses to characterize the extent and distribution of extant genetic diversity.

**Figure 1.**
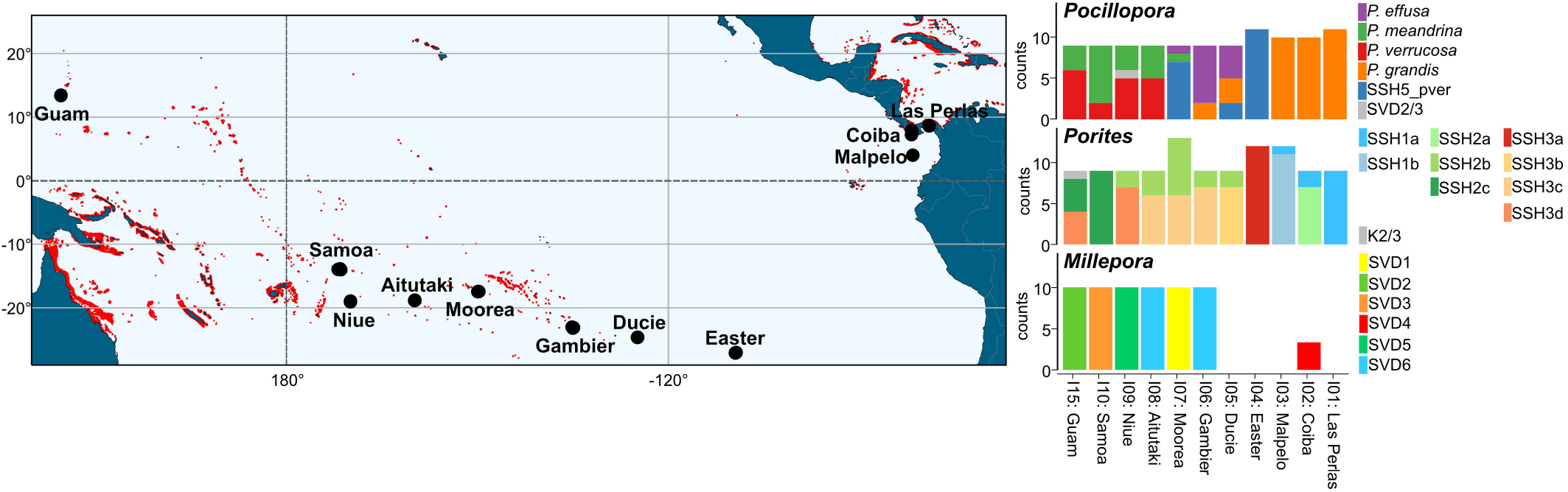
Biogeography of genetic delineations of the genera *Pocillopora, Porites*, and *Millepora* across their Pacific spread. The number of samples belonging to a given genetic delineation are given for each of the genera. Reference reefs from the United Nations Environment Programme—World Conservation Monitoring Center’s (UNEP-WCMP) Global Distribution of Coral Reefs dataset (Spalding et al. 2001; Wri 2005) are plotted in red using reefMapMaker (Hume and Voolstra 2022).

### Genetic delineation and biogeography of *Pocillopora* spp

For *Pocillopora*, Maximum Likelihood (ML; Supplementary Figure 1) and SVDquartet (used to estimate the species-tree topology based on SNP data under a multi-species coalescent model; Figure 2A) trees shared a similar topology, with five well supported clades (SVD1-SVD5) containing the same constituent samples and with the same single outlier sample (I09S03C010). The SVD1 clade was more distantly related to the other SVD clades and we could observe two pairs of sister clades: SVD3 & SVD5 and SVD2 & SVD4. These clades were further supported through estimation of individual ancestry coefficients using sNMF genetic clustering and principal component analysis (PCA), which predicted five ancestral populations (K1-K5). While minimal introgression was apparent across most samples, several subsets of samples from specific islands, and resolving to specific subclades of the SVD tree, displayed higher levels of introgression (e.g., K3_I15, K4_I05, and K5_I07; Figure 2C). The one sample that grouped as an outlier in the ML and SVDquartet trees demonstrated a hybrid ancestry with an approximately 50/50 split of the ancestral populations otherwise associated with samples from the SVD2 and SVD3 groupings. Despite the island-specific patterns of admixture in some lineages, sNMF analysis within the identified SVD lineages identified no further genetic clustering. The SVD groupings were recapitulated in the PCA across the four highest scoring principal components (PCs; Figure 2B). We tested multiple SSH by coalescence analysis using two replicate runs of BFD* with samples categorized by SVD group. The most likely species hypotheses differed between runs, but the second most likely hypothesis in both runs designated each SVD a separate species (Supplementary Table 2). We adopted this 5-species SSH as our working hypothesis. Of note, the difficulty in such assignments lies in avoiding mixing of isolated populations and species. Given that one has to drastically reduce sampling size for this type of analysis, we chose to discard SSH whose likelihood ranks vary between replicate runs. To assign species names to our SSH, two representatives for each SVD grouping were mapped onto the *Pocillopora* reference sequences from (Oury et al. 2022) to then call the species-diagnostic SNPs generated therein. While SVD1-SVD4 could be assigned to the following species names: *P. effusa* (SVD1), *P. meandrina* (SVD2), *P. verrucosa* (SVD3), and *P. grandis* (SVD4), SVD5 was most similar to a lineage with no formal name that is most closely related to *P. verrucosa* (hereafter referred to as SSH5_pver; Supplementary Figure 2). When assessing how samples from the different species were distributed across our sampling regime, we found that species exhibited region-specific distributions with each region composed of adjacent islands (Figure 1). *Pocillopora meandrina* and *P. verrucosa* were distributed in the western Pacific, *P. effusa* and SSH5_pver in the central Pacific, and *P. grandis* in the eastern and central Pacific. At all but the four most eastern sites (Easter, Malpelo, Coiba, and Las Perlas), multiple species were found in sympatry (Figure 1). Notably however, *P. verrucosa* and SSH5_pver as well as *P. meandrina* and *P. grandis*, the two most genetically similar species pairings, were only found in allopatry.

**Figure 2.**
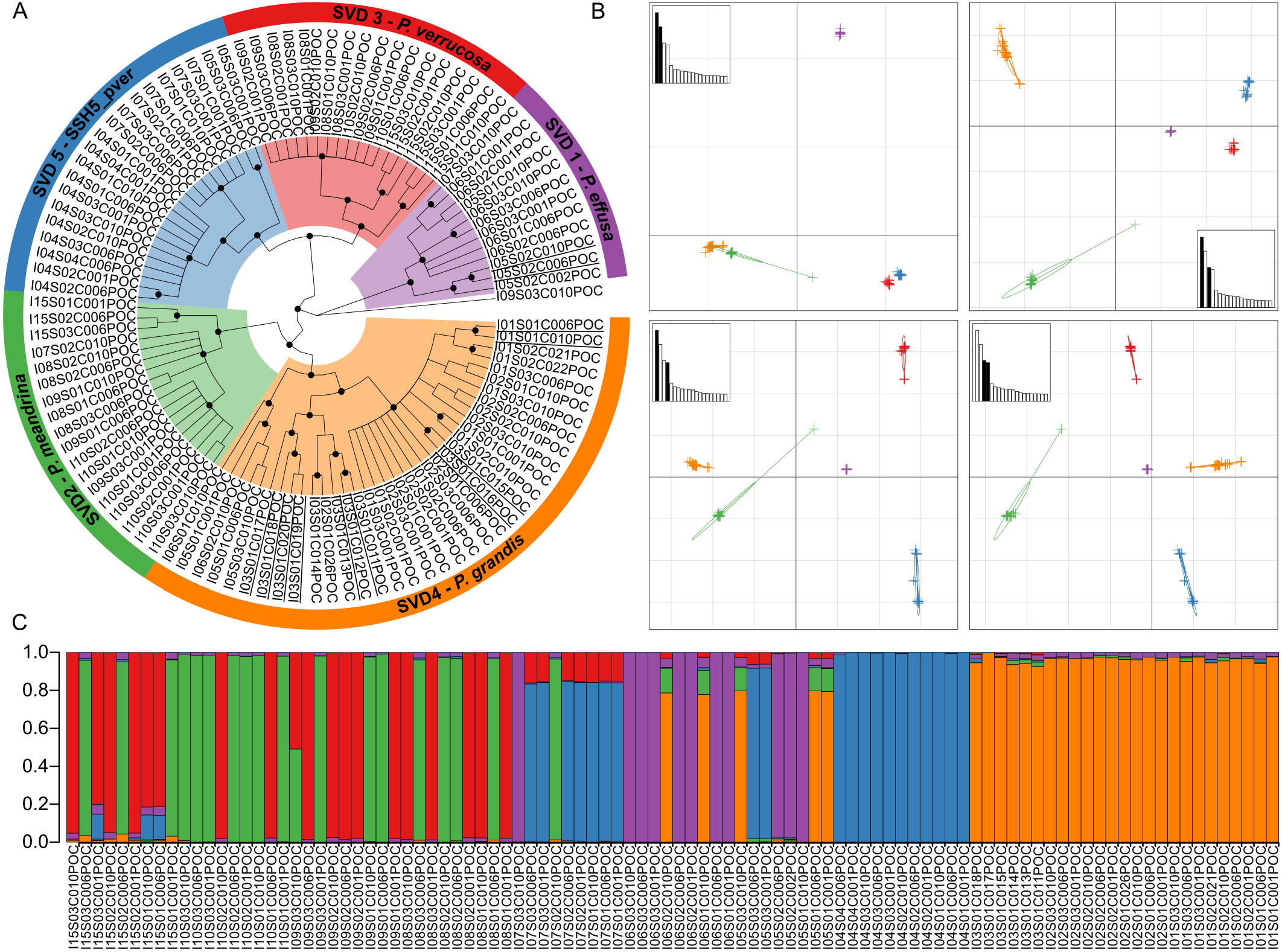
Genetic delineation in *Pocillopora*. A) SVDquartet tree. Node support >= 80% from 100 bootstraps is indicated with a black dot. Samples are grouped into five clades (SVD1-SVD5) based on tree morphology and sNMF analysis which form the secondary species hypotheses (SSH). Species names resulting from subsequent analyses (see relevant section of this study) have been annotated. Samples not included in the sNMF analysis (due to removal of clonal/multilocus lineages) are underlined. B) Principal component analysis (PCA) based on genome wide SNPs. The relative eigen values are shown in the subplots with the displayed eigenvectors shaded black. C) Hierarchical genetic clustering using sNMF. Bars in each column represent the contribution of predicted ancestral lineages for each sample. Samples are ordered longitudinally by island.

### Genetic delineation and biogeography of *Porites* spp

The ML (Supplementary Figure 1) and SVDquartet (Figure 3A) trees generated for *Porites* both supported the existence of three clades (SVD1-SVD3) with SVD2 and SVD3 more closely related to each other than SVD1. All samples resolved within one of these clades except for sample I15S02C011 that resolved between SVD1 and the other two clades. These clades were further supported by sNMF genetic clustering and principal component analysis (PCA). The sNMF analysis suggested three ancestral populations (K1-K3) with minimal introgression apparent (Figure 3C). Notably, there was exact agreement between SVDquartet clade placement and sNMF assignments (according to highest admixture coefficient; hereafter, SVD1 will be related to K1, SVD2∼K2 etc.) for all but three of the samples, which exhibited significant levels of admixture (see Supplementary Table 3). Unlike in *Pocillopora*, genetic subclustering was detected within each of the sNMF groupings with two, three, and four ancestral populations predicted for K1, K2, and K3 (hereafter referred to as K1a, K1b, K2a etc.; Figure 3C), respectively. These subclusterings matched the subclade structure in the SVD groupings (exact match in SVD1 and SVD3; close match in SVD2; Figure 3A) and largely demonstrated island-specific, but not endemic, patterns (Figure 1). We tested the SSH by coalescence analysis using the sNMF subcluster groupings (i.e., K1a, K1b, K2a etc.). As in *Pocillopora*, the most likely species hypotheses differed between runs, but the second most likely hypothesis in both runs designated K1, K2 and K3 as distinct species. We adopted this 3-species SSH as our working hypothesis and hereafter refer to SSH1-SSH3 in reference to K1-K3 (and SSH1a in reference to K1a, etc.). Similar to *Pocillopora*, each of the SSH were distributed across multiple islands with one or two lineages found at each island. SSH2 and SSH3 were most widely distributed with each found in seven islands (absent in only Malpelo and Las Perlas), while SSH1 was found only in the easternmost three sites (Malpelo, Coiba, and Las Perlas; side-by-side with SSH2; Figure 1). In contrast to *Pocillopora* where the genetically similar species pairings were found in allopatry, the more genetically similar species, SSH2 and SSH3, were found in widespread sympatry.

**Figure 3.**
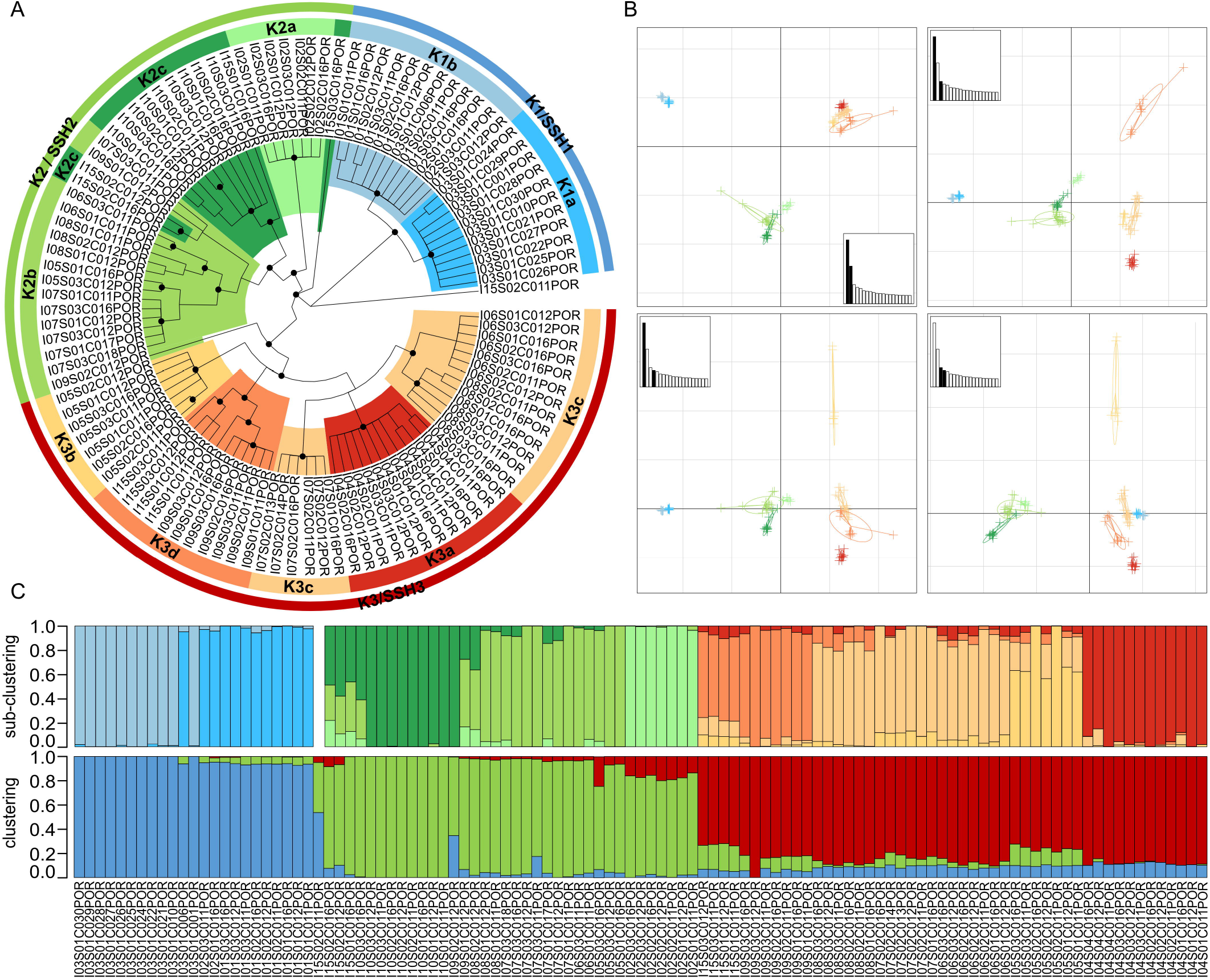
Genetic delineation in *Porites*. A) SVDquartet tree. Node support >= 80% from 100 bootstraps is indicated with a black dot. Samples are grouped into three main clades (K1-K3) and two, three and four subclades, respectively (K1a, K1b etc.) based on tree morphology and sNMF analysis with the main clades representing the three secondary species hypotheses (SSH; annotated). B) Principal component analysis (PCA) based on genome wide SNPs. The relative eigenvalues are shown in the subplots with the displayed eigenvectors shaded black. C) Hierarchical genetic clustering using sNMF across all samples with main clades below and subclades above. Bars in each column represent the contribution of predicted ancestral lineages for each sample. Samples are ordered by cluster and then longitudinally by island.

### Genetic delineation and biogeography of *Millepora* spp

In *Millepora*, agreement in topology between the ML (Supplementary Figure 1) and SVDquartet (Figure 4A) trees was poor, unlike in *Pocillopora* and *Porites*. While the SVD tree showed relatively strong bootstrap values that supported the existence of six clades (SVD1-SVD6), the branches of the ML tree lacked support, and distinct clades were difficult to resolve with the exception of the *M. intricata* samples (identified as a distinct species based on field-based morphological assessment; see Methods) that were resolved separately from all other samples in both trees. The sNMF analysis predicted four ancestral populations (K1-K4) with levels of introgression varying considerably. Unlike in *Pocillopora* and *Porites*, the sNMF ancestral populations did not correspond directly to the SVD groupings, and rather, discrete combinations of the four ancestral coefficients resolved six groups that corresponded to the SVD clades (Figure 4C). PC1 of the PCA analysis grouped the *M. intricata* samples separately from all others, while the remaining samples grouped according to the five remaining SVD groups across PCs 2-4 (Figure 4B). Due to the low levels of support in the ML tree, the higher levels of introgression from predicted ancestral populations in extant samples (sNMF analysis), and the lack of direct correspondence between the sNMF ancestral populations and the SVD groupings, we considered the five non-*M. intricata* SVD groupings (SVD2-SVD6) to be representative of a single species, designated as *Millepora* cf. *platyphylla* (based on Boissin et al. 2020). Unlike in *Pocillopora* and *Porites, Millepora* SVD clades were all endemic to specific islands except for SVD6. Nevertheless, a split at the basal node of the SVD6 group partitioned the samples into two clades with all members endemic to either Gambier or Aitutaki (Figure 1, Figure 4A).

**Figure 4.**
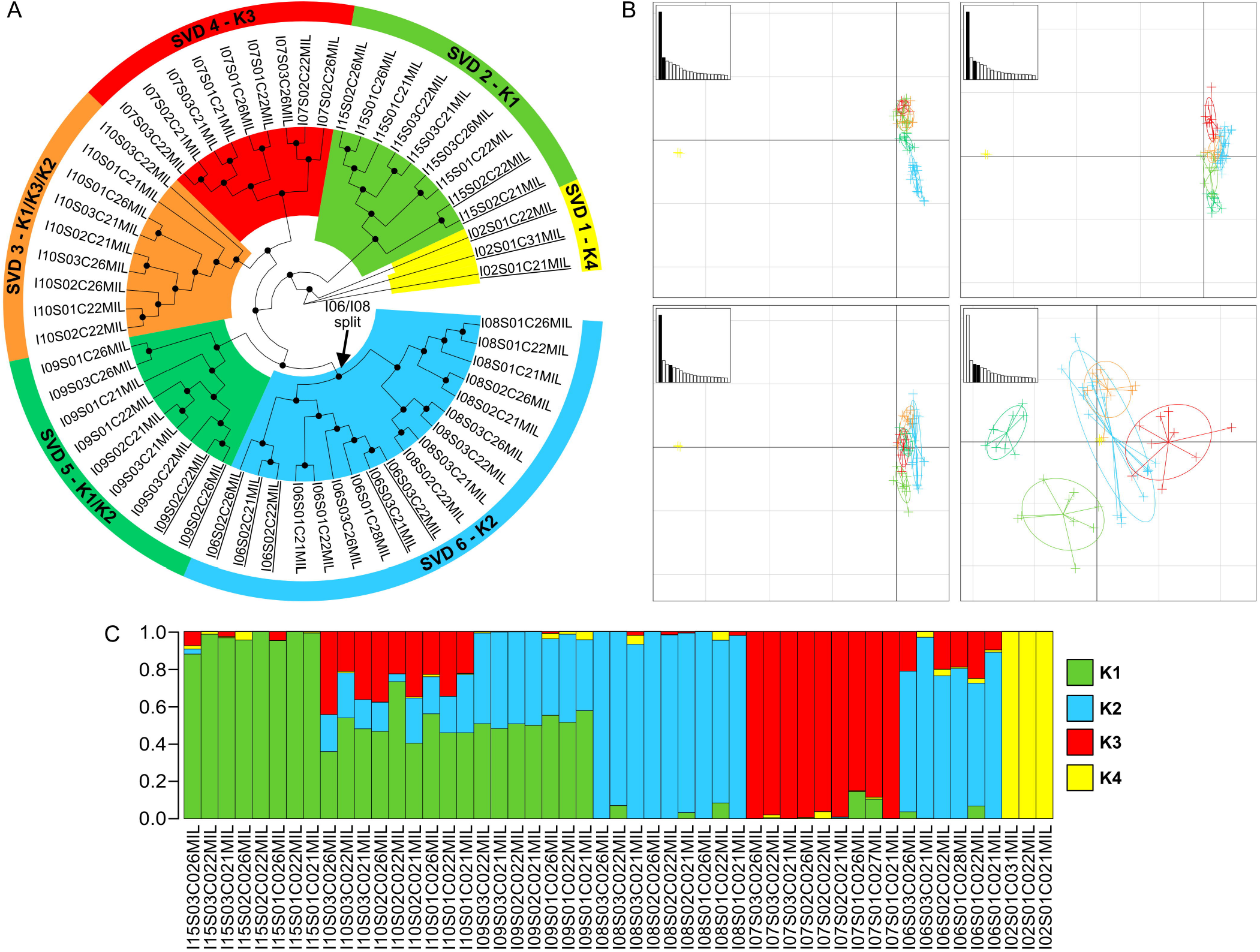
Genetic delineation in *Millepora*. A) SVDquartet tree. Node support >= 80% from 100 bootstraps is indicated with a black dot. Samples are grouped into six clades (SVD1-SVD6) based on tree morphology and sNMF analysis. Samples not included in the sNMF analysis (due to removal of clonal/multilocus lineages) are underlined. B) Principal component analysis (PCA) based on transcriptome wide SNPs. The relative eigen values are shown in the subplots with the displayed eigenvectors shaded black. C) Hierarchical genetic clustering using sNMF. Bars in each row represent the contribution of predicted ancestral lineages for each sample. Samples are ordered longitudinally by island.

### Morphological analysis

In parallel to the genetic analyses, we also conducted morphological analyses that employed a multivariate suite of macromorphological measurements (see Methods). This served a dual purpose: first, we wanted to assess whether we could identify morphological variation between populations, and second, we wanted to assert whether such variation corresponded to the genetically identified SSH, to provide genetic and morphological evidence of the species designations. Morphological analysis revealed structured variation in *Pocillopora* (Supplementary Figure 3), *Porites* (Supplementary Figure 4), and *Millepora* (Supplementary Figure 5). We also found significant PERMANOVA results when testing the multivariate morphological characteristics against the genetic delineations, which demonstrated that some of the variation could be explained by the SSH designations (*Pocillopora*: p = 0.013, *Porites*: p = 0.001, and *Millepora*: p = 0.003). Critically however, for each of the coral genera, there was no combination of morphological features that could identify the genetic resolutions; rather, each of the morphotype categorizations contained multiple SSH. Within-genus pairwise PERMANOVA testing identified differences between all pairings except for *P. meandrina* vs. *P. verrucosa* and *P. meandrina* vs. SSH5_pver in *Pocillopora*, and SSH6 vs. SSH5, SSH6 vs. SSH2, SSH4 vs. SSH3, SSH4 vs. SSH2, SSH5 vs. SSH3, and SSH3 vs. SSH2 in *Millepora*. All within-*Porites* pairings returned significant results.

### Evolutionary history of among- and within-species genetic differentiation

To further resolve evolutionary history in the *Pocillopora* and *Porites* species designations, we investigated introgression and pairwise divergence for the SSH (Figure 5). For *Millepora*, an insufficient number of SNPs and representative samples meant that these analyses were not conducted. In *Pocillopora*, three admixture events were predicted and verified based on significant results in the relevant *f*_4_ indices (Reich et al. 2009; Figure 5B&C) where two of the admixture events were from ancestral nodes to child clades and the third was inter-SSH from *P. verrucosa*_I06 to *P. meandrina*_I09. The resulting TreeMix consensus tree showed a similar topology to that of the SVD tree with those groups from *P. effusa* more differentiated from the other SVD groups and with two pairs of sister clades formed by *P. meandrina & P. grandis* and *P. verrucosa &* SSH5_pver (Figure 5B). After accounting for the three introgression events, residuals (obtained from fitting the tree model to the observed data) indicated potential remaining admixture within each of the five species (Figure 5A). Similar to the Treemix tree, the f_2_ values (Supplementary Figure 6A) and the distributions of genomic Weir’s *F*_ST_ values computed from 500 bp sliding windows (Supplementary Figure 6B) recapitulated the high divergence of *P. effusa* from the other more genetically similar two species pairs (*P. meandrina & P. grandis* and *P. verrucosa &* SSH5_pver). Taken together, these findings reinforce the pattern in *Pocillopora* that the genetically closely related sister clade species pairings (*P. meandrina & P. grandis and P. verrucosa &* SSH5_pver) are found in allopatry rather than sympatry. In *Porites*, three admixture events were predicted and confirmed (f_4_ significant results; Figure 5D&E) with one within-SSH, one inter-SSH, and one from an ancestral to a child clade. Similar to *Pocillopora*, the resulting TreeMix consensus tree topology agreed with that of the SVD tree, with SSH1 being more differentiated compared to SSH2 and SSH3. After accounting for the three admixture events, residuals indicated potential remaining admixture within each of the three SSH and also between SSH2 and SSH3 (Figure 5D). The f_2_ values (Supplementary Figure 6C) and the distributions of Weir’s *F*_ST_ from 500 bp sliding windows (Supplementary Figure 6D) both supported the relative divergence of SSH1 from the other two SSH, and in contrast to *Pocillopora*, it also showed a correlation between sympatry and genetic relatedness.

**Figure 5.**
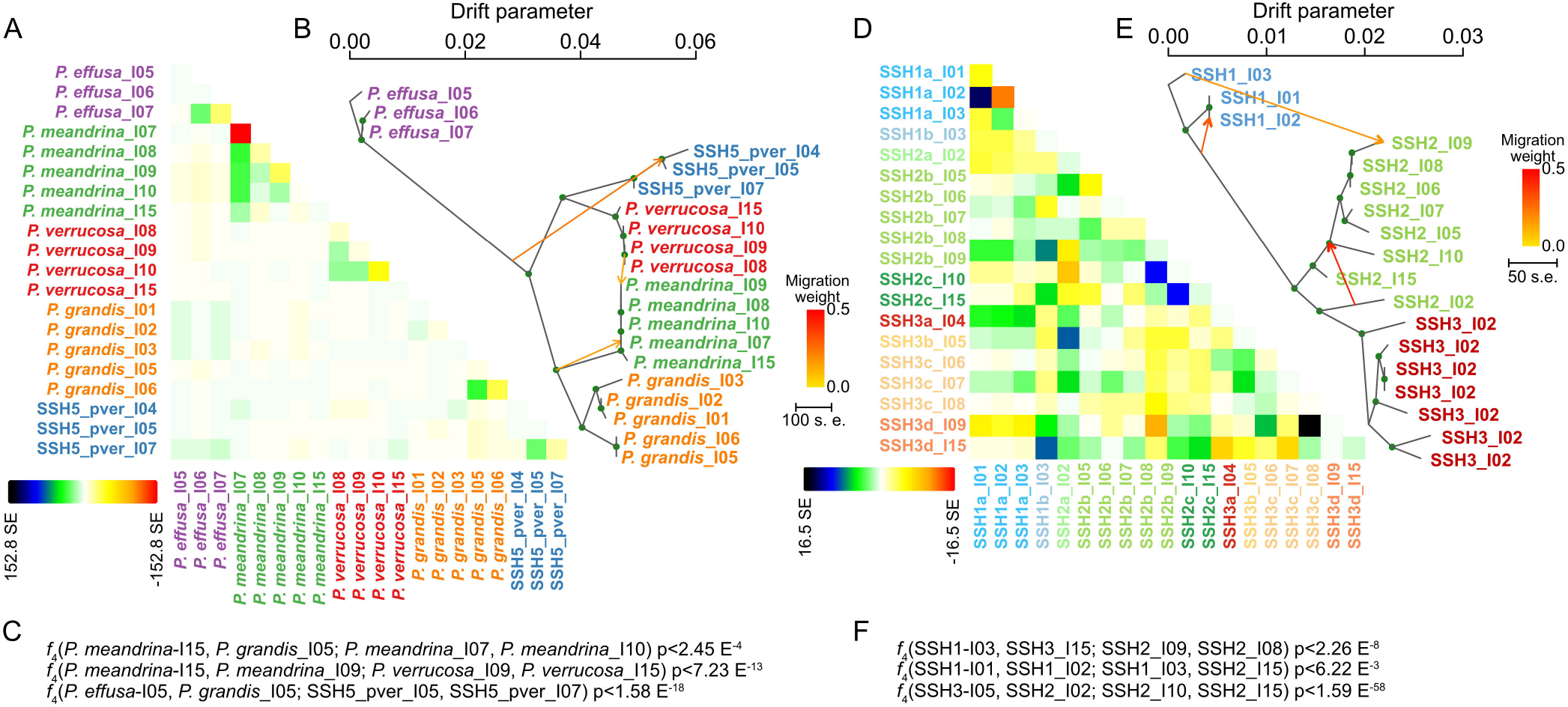
TreeMix consensus tree with migration events and residuals for *Pocillopora* and *Porites*. A&D) residuals indicating potential remaining admixture between samples in *Pocillopora* and *Porites*, respectively, after accounting for the three migration events annotated in the TreeMix tree. B&E) Treemix consensus tree with migration events and their weights annotated. C&F) Corresponding *f*_4_ indices.

### Correlation of genetic relatedness with geographic distance and climate

To assess for isolation-by-distance and a possible effect of temperature regime on the evolutionary trajectories of the species, we separately tested for correlation between genetic distance (encoded as pairwise *F*_ST_/(1-*F*_ST_) distances generated from pairwise group *F*_ST_ distances; see Methods) and both geographical distance (calculated as over-water distance between islands) and climate (represented by historical temperature difference based on island-wise Euclidean historical temperature distances generated from a reduced dimensionality representation; see Methods and Lombard et al. 2022).

In *Pocillopora*, separate correlation analyses of genetic distance against geographical distance and temperature difference returned non-significant results across- and within-species (Supplementary Table 4). By contrast, significant results were returned for *Porites* across all species (geography, r=0.397, p=0.005; temperature, r=0.361, p=0.020) and within SSH2 (geography, r=0.585, p=0.036; temperature, r=0.641, p=0.010) and SSH3 (geography only, r=0.591, p=0.008). In *Millepora*, a significant result was observed for all lineages against temperature r=0.930, p=0.019) but not geography (r=0.722, p=0.053). For each coral genus, geographical distance significantly correlated with historical temperature distance (Supplementary Table 4).

### Influence of temperature regime on evolutionary trajectories

To investigate whether temperature (as a prevalent stressor of the coral metaorganism) was a potential driver of species divergence pattern in *Pocillopora* and *Porites*, we assessed whether SNPs that aligned with historical temperature regimes exist. Indeed, we could identify SNPs in *Pocillopora* and *Porites* whose local allelic frequencies could be predicted by historical temperature regime (i.e., temperature outlier SNPs; Pocillopora – 38,229 out of 461,989 SNPs (8.27%); Porites – 6,441 out of 57,966 SNPs (11.11%); at q value <0.1 from the unliked dataset, see Methods; Figure 6). To determine which of these SNPs were potentially involved in species divergence patterns we queried which of the SNPs originated from a set of genomic islands of differentiation (GID) defined as the 1% of the 500 bp bins with the highest mean *F*_ST_ value among SSH for *Pocillopora* and *Porites*. In *Pocillopora*, 20 of the top 100 temperature outlier SNPs (ranked by p value) were located in these GIDs (Figure 6). By contrast, *Porites* had only 3 such SNPs in GIDs, suggesting that species differentiation is more closely correlated to thermal history in *Pocillopora* than in *Porites*. We also investigated whether this difference could be due to drift or actively maintained due to divergent adaptation. In *Pocillopora*, 21% of the whole genome was predicted to be under divergent selection (at the level of between-SSH comparisons), a considerably higher proportion than the 7% predicted in *Porites*. However, independent of this baseline difference between the two genera, in *Pocillopora* 41% of SNPs (41 out of 100) in the top 100 temperature outliers were predicted to be under selection, and this proportion grew to 61% (12 out of 20) when considering those outlier SNPs under selection and residing in GIDs, approximately double and triple that of the proportion of SNPs under divergent selection across the whole genome (21%). By contrast, in *Porites* a decrease from the 7% of SNPs under divergent selection across the whole genome was observed when considering either those outlier SNPs predicted to be under selection (2%; 2 out of 100) or those under selection and residing in GIDs (0%; 0 out of 3). This suggests that while the differentiation and distribution of the *Pocillopora* species are due to and/or maintained by divergent selection driven by temperature regimes, in *Porites* divergence among SSH is likely due to either drift or other environmental factors (of note, many environmental factors correlate with temperature).

**Figure 6.**
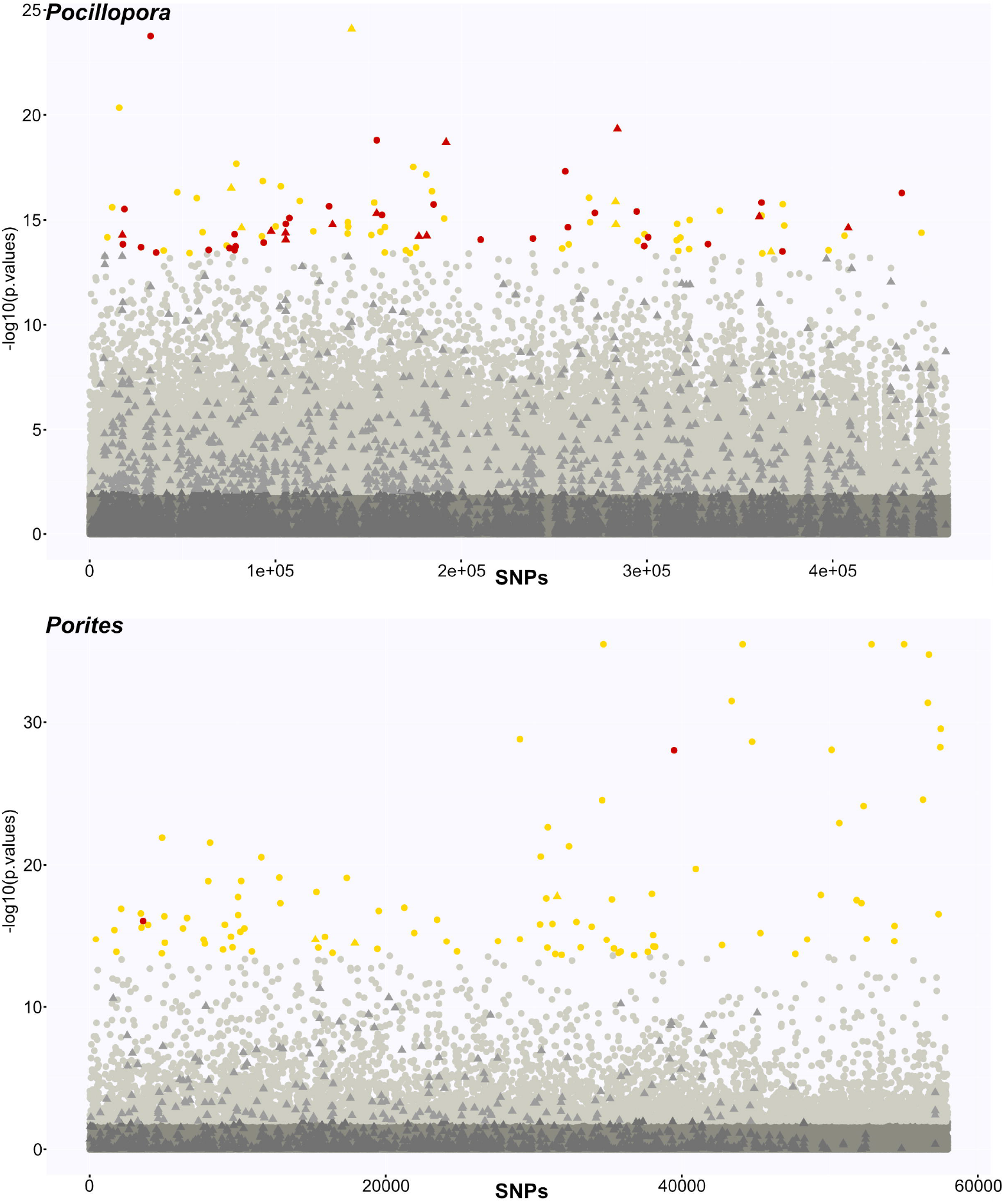
Manhattan plot displaying the association to historical temperature variation of the linkage disequilibrium-filtered genome-wide SNPs computed by RDA analysis. The x axis gives cumulative position in the genome with contigs sorted by decreasing size order. p values used to generate the y axis are from the RDA analysis. Circles: SNPs not included in genomic islands of differentiation between SSH (GID); triangles: SNPs included in GIDs; dark gray: SNPs not significantly associated to historical temperature variation; light gray: SNPs significantly associated to temperature variation; gold: top 100 temperature outlier SNPs; red: SNPs among the 100 top temperature outliers under divergent selection among species according to Flink analysis.

## DISCUSSION

Our primary species hypothesis of one species per coral genus, based on *in situ* morphological assessment, was genetically validated only for *Millepora* cf. *platyphylla*. The corals collected and putatively designated as *Pocillopora meandrina* and *Porites lobata* produced SSH of 5 and 3 species, respectively, that were supported by multiple genetic analyses (Figures 2, 3, 4 & 5; Supplementary Figure 6). Subsequent morphological analyses based on underwater colony pictures of the corals resolved a number of morphotypes (Supplementary Figures 3-5) that showed correlation with the genetically resolved lineages in all three genera. However, multiple genetic lineages were associated with all designated morphotypes and the identification of lineage-diagnostic morphological features was not possible based on underwater colony pictures. Our finding underlines the difficulty of defining morphological criteria that can be readily derived from photographs and therefore supports the well-established existence of high morphological plasticity in corals, as exemplified by the genus *Pocillopora* (Erika C. Johnston et al. 2017; Marti-Puig et al. 2014; Paz-García et al. 2015). Further, it reinforces that plasticity is a confounding factor to achieve effective taxonomic resolutions (Erika C. Johnston et al. 2017; Todd 2008) and identification of species in the field. In the context of conservation strategies, the disparity between morphology- and genetics-based species resolution highlights the necessity for robust molecular characterization to be able to resolve species diversity.

The three genera demonstrated very different degrees of genetic diversity and connectivity despite being sampled across the same environmental range and same sites. This has implications for informing conservation strategies. Our genetic analyses suggest that *Pocillopora* is characterized by a higher level of within-species connectivity than *Porites*. A correlation between genetic dissimilarity and geographic distance as well as within-species genetic structuring (by sNMF genetic clustering; Figures 1&3) was only detected in *Porites* (in line with less resolutive past studies; Boulay et al. 2014; Baums, Boulay, and Polato 2012), and residuals (after TreeMix consensus tree generation) gave a considerably stronger within-species introgression signal in *Pocillopora* than in *Porites* (Figure 5). Maintenance of connectivity to ensure exchange of alleles is a key aim of resilience-based conservation management strategy (Mcleod et al. 2019) and has been demonstrated to facilitate robustness and recovery to stress in reef ecosystems (Palumbi 2003; Van Oppen and Gates 2006; Colton et al. 2022). Therefore, whilst the higher number of species resolved in *Pocillopora* may suggest a greater conservation effort is required (greater species diversity to protect), its higher connectivity, in concert with its relatively high levels of species sympatry at certain sites (all species are present when considering only two sites, e.g. I05 - Ducie and I08 - Aitutaki; see also E. C. Johnston et al. 2022), mean that core levels of diversity protection and resilience maintenance may be accomplished with a relatively focused effort. However, whether observed species are truly adapted to the given location or whether their presence is the result of / dependent on migration from a seed site (see below) must be carefully considered. In contrast to *Pocillopora*, the lower levels of connectivity and relatedly higher levels of genetic structure in *Porites* suggest a greater spatial area of protection would be required to conserve genetic diversity, and promote resilience, especially considering the geographic isolation of SSH1 in the eastern Pacific (I01-03; Las Perlas, Coiba, and Malpelo). While *Pocillopora* and *Porites* resolved into multiple distinct species, *Millepora* was resolved as a single species with highly endemic populations containing varying levels of admixture. In the context of maintaining genetic diversity, the *Millepora* population of I10 (Samoa) would be prioritized for protection over others as this population has the highest proportion of admixture encompassing all three of the predicted ancestral populations. A recent study by (Buitrago-Lopez et al. 2022) also detected distinct differences in connectivity between two reef-building species of the family Pocilloporidae that they posited may, at least partially, be due to different reproductive modes (broadcast spawner, high connectivity, low genetic structure; brooder, low connectivity, high genetic structure). Reproductive mode could explain the differences in connectivity we observe here but has not been characterized for the corals sampled. Further, generalizations based on other members of the respective genera are difficult, as *Pocillopora* and *Porites* both contain species that broadcast spawn and internally fertilize (brood), with some species exhibiting location-dependent reproductive modes (Brown et al. 2020; Richmond and Hunter 1990; Fadlallah 1985; Sier and Olive 1994; Stimson 1978; Schmidt-Roach et al. 2012). Independent of identifying a causal factor for differences in connectivity, it is clear that species- and even population-specific characterizations of connectivity are a necessity in designing effective strategies (and conservation areas/zones) for genetic diversity maintenance.

Our dataset comprising samples of multiple species of *Pocillopora* and *Porites* across the same basin-wide range has enabled us to demonstrate contrasting evolutionary histories in these corals, despite exposure to the same past thermal, and arguably environmental, regimes. However, our finding that climate plays a greater role in species divergence in *Pocillopora* than in *Porites* contrasts the non-significant (*Pocillopora*) and significant (*Porites*) correlations between genetic distance and historical thermal distances (Supplementary Table 4) detected in the genera. Given the significant correlation to thermal distance in *Porites*, one might have expected historical thermal variation to play a role in *Porites* speciation/species divergence. Rather, it would appear that the significant correlation to temperature distance is the product of correlation between the geographic and temperature distances (r=0.564, p=0.000, Supplementary Table 4) with the significant genetic and geographic correlation driven by the relatively low dispersal of *Porites*. In *Pocillopora*, the lack of correlation between genetic distance and thermal distance is likely a product of the higher overall species sympatry observed in *Pocillopora* (Figure 1) that may be explained by high levels of dispersion and subsequent migration. Migration would enable a continuous influx of individuals with minimal genetic differentiation from seed sites to destination sites. These individuals would be adapted to the seed sites and may not survive in the destination sites over the time frames required for selection and genomic adaptation to the destination sites to act. In this scenario, individuals from a given species may be distributed across a wide range of sites while remaining adapted only to the original seed environment. This would concurrently explain the observed high overall species sympatry, lack of genetic distance correlations, yet the considerable effect of historical temperature variation on the maintenance of species divergence (if not on the speciation process itself) in *Pocillopora*. The larger effect of historical temperature variation on the evolutionary histories of the *Pocillopora* species, which is generally recognised as a stress-sensitive genus (Darling et al. 2012), could result from a higher exposure to environmental perturbations at the reef spatial scale. This is because it generally inhabits more variable and higher energy environments on the reef than *Porites* which is more broadly distributed (Veron 2000). It could also be symptomatic of a specialization/niche adaptation strategy (as opposed to being a more environment generalist like *Porites*) whereby any perturbation from the niche environment would necessitate an adaptive response to survive due to a limited acclimative potential (Ziegler et al. 2014). However, this explanation is less parsimonious given the acknowledged high phenotypic plasticity of the genus considered to be a driving factor in its evolutionary success (Erika C. Johnston et al. 2017). By contrast, the relative lack of divergent selection among *Porites* species suggests that temperature perturbations did not influence the speciation process in this genus, in the sense that the elicited adaptive responses (resulting in at least some of the temperature outlier SNPs revealed by the genotype/environment association analyses) did not participate to the species differentiation. It could also be argued that the adaptive potential or *Porites* is limited, but this is unlikely given the globally recognised resilience of this genus (Darling et al. 2012; Sawall et al. 2015; Ziegler et al. 2014). The genomic data produced here can provide the starting point for future functional analyses investigating the genes and pathways associated with the temperature outlier and GID SNPs identified, thus enabling cellular insights into adaptation potential and its underlying mechanisms.

Contemporary resilience-based conservation strategies do not fully consider how corals may adapt to future challenges. Given the differences between species found across the same distribution here, it is interesting to speculate on species being more or less successful under climate change. *Pocillopora* species appear more linked to the environment than *Porites* and therefore likely display a narrower set of physiological tolerances. However, *Pocillopora* is known to be a speciose genera (Pinzón et al. 2013) and while some of those species may be lost with the continuing and growing climatic perturbations predicted in the coming decades, surviving genotypes will have been selected for the future environments. Coupled with the high dispersion capability of the genus - granting access to the widest possible range of habitats - this may result in a relatively successful strategy for the *Pocillopora* genus, enabling recolonization of future reefs by its more stress tolerant species. Conservation efforts applied to such speciose, niche-adapted, and high-dispersal coral genera may catalyze the recovery of reefs by identifying those genotypes most likely to survive future perturbations, and where possible, protecting them from further non-climatic perturbations while facilitating their already high dispersal capacities to ensure the seeding of as wide a range of reef ecosystems as possible. In contrast to the more niche-adapted corals, generalist corals such as the *Porites* species identified in this study may respond differently to future perturbations. So long as the magnitude of perturbations are within the buffering capacity of their wider physiological tolerances, the generalist strategy of these corals may be successful in the long-term. However, as these buffering capacities are surpassed, the lower dispersal capacity of these species will mean a reduced ability to recolonize alternative environments and the genera may suffer losses. In contrast to the targeted strategy of the niche-adapted corals, conservation of generalist corals should likely focus on mitigating local aggravating stressors (e.g., managing pollutant and nutrient input, limiting physical disruption due to coastal development, or preventing overfishing) to maximize the buffering capacity of the corals across as many of the available genotypes as possible.

## ACKNOWLEDGEMENTS

Special thanks to the *Tara* Ocean Foundation, the R/V *Tara* crew and the *Tara* Pacific Expedition Participants (https://doi.org/10.5281/zenodo.377776). We are keen to thank the commitment of the people and the following institutions for their financial and scientific support that made *Tara* Pacific expedition possible: CNRS, PSL, CSM, EPHE, Genoscope/CEA, Inserm, Université Côte d’Azur, ANR, and the Tara Ocean Foundation’s teams, crew, President Etienne Bourgois and its partners agnès b., UNESCO-IOC, the Veolia Environment Foundation, Région Bretagne, Billerudkorsnas, Amerisource Bergen Company, Lorient Agglomeration, Smilewave, Oceans by Disney, the Prince Albert II de Monaco Foundation, L’Oréal, Biotherm, France Collectivités, Fonds Français pour l’Environnement Mondial (FFEM), the Ministère des Affaires Européennes et Etrangères, the Museum National d’Histoire Naturelle. *Tara* Pacific would not exist without the continuous support of the participating institutes. The authors also particularly thank Serge Planes, Denis Allemand and the *Tara* Pacific consortium. This is publication number 25 of the *Tara* Pacific Consortium

The acknowledgements to local people and authorities who made this study possible can be found in the associated Zenodo deposition (Hume et al. 2022). We would like to thank Hans-Joachim Ruscheweyh as well for lending us computing time at ETH for some of the BFD* replicate runs. Work from BP, ER, DF is supported by the French Government (National Research Agency, ANR) through the grant “Coralgene” ANR-17-CE02-0020 as well as the “Investments for the Future” programs LABEX SIGNALIFE ANR-11-LABX-0028 and IDEX UCAJedi ANR-15-IDEX-0001. The *Tara* Pacific expedition would also not have been possible without the participation and commitment of over 200 scientists, sailors, artists and citizens (see https://zenodo.org/record/3777760#.YfEEsfXMLjB).

## SUPPLEMENTARY ITEMS

### SUPPLEMENTARY FIGURE LEGENDS

**Supplementary Figure 1.**
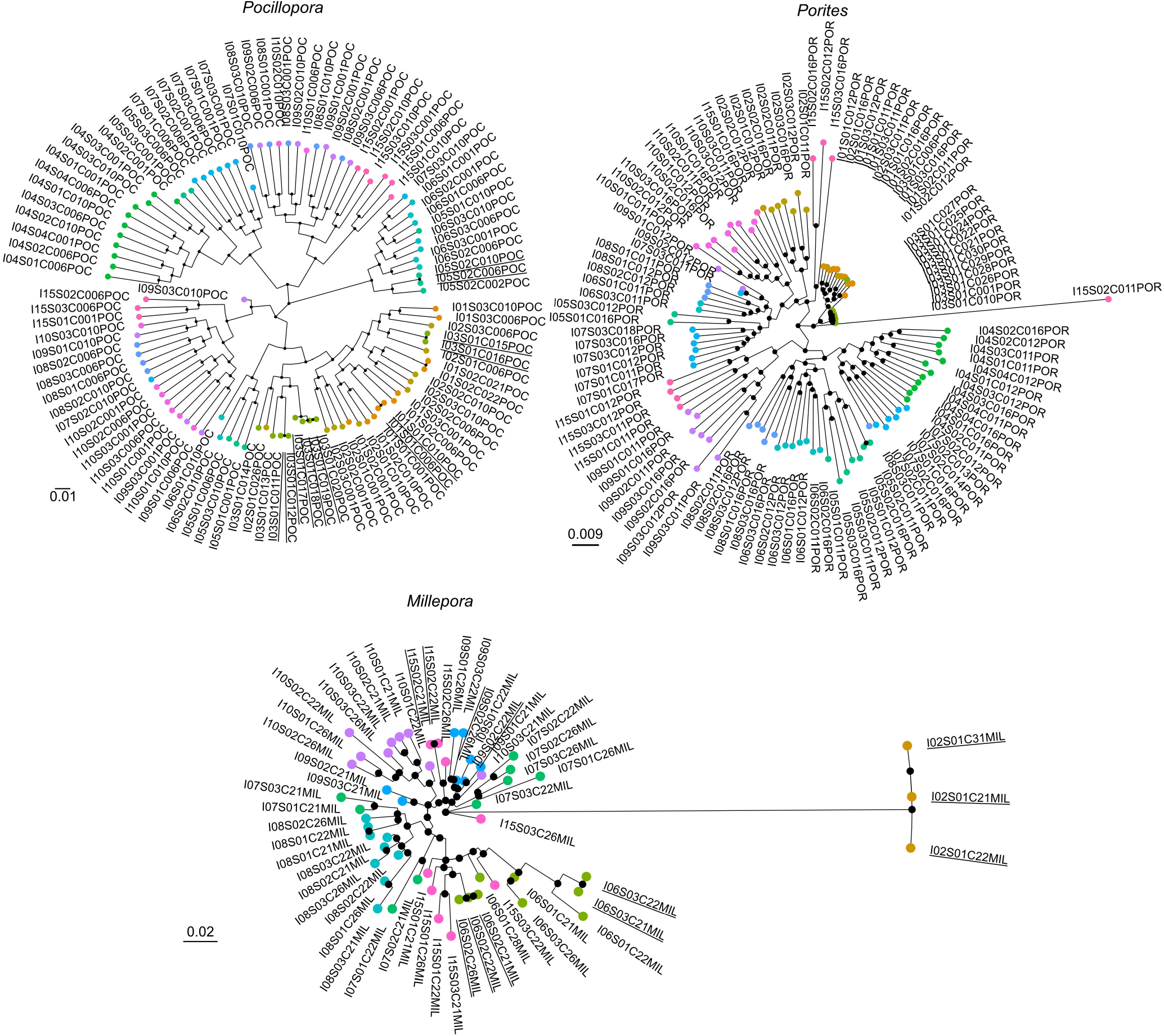
Maximum likelihood phylogenetic trees for *Pocillopora, Porites*, and *Millepora*. Node support ≥ 80% is indicated with black circles. Leaves are colored according to the island of collection.

**Supplementary Figure 2.**
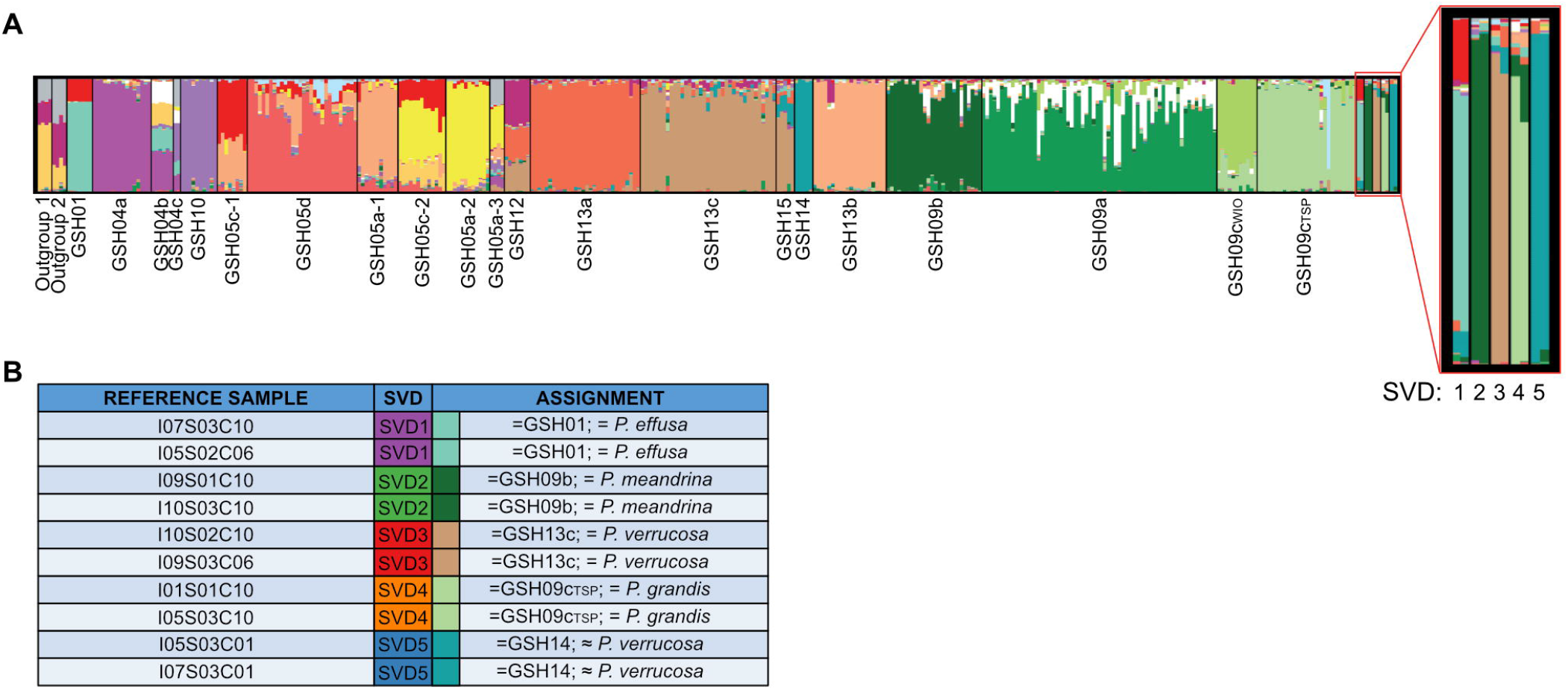
Assignment of taxonomy in *Pocillopora*. A) sNMF results from the inclusion of two representative samples for each SVD lineage with the dataset of (Oury et al. 2022). Bars show the contribution of each of 20 ancestral lineages to the samples. The ten samples from this study are on the right of the figure. Samples are grouped according to the identified lineages. B) Assignation of taxonomy to the ten reference samples. Colors correspond to the SVD lineages used in the main manuscript and the sNMF ancestral lineages used in A.

**Supplementary Figure 3.**
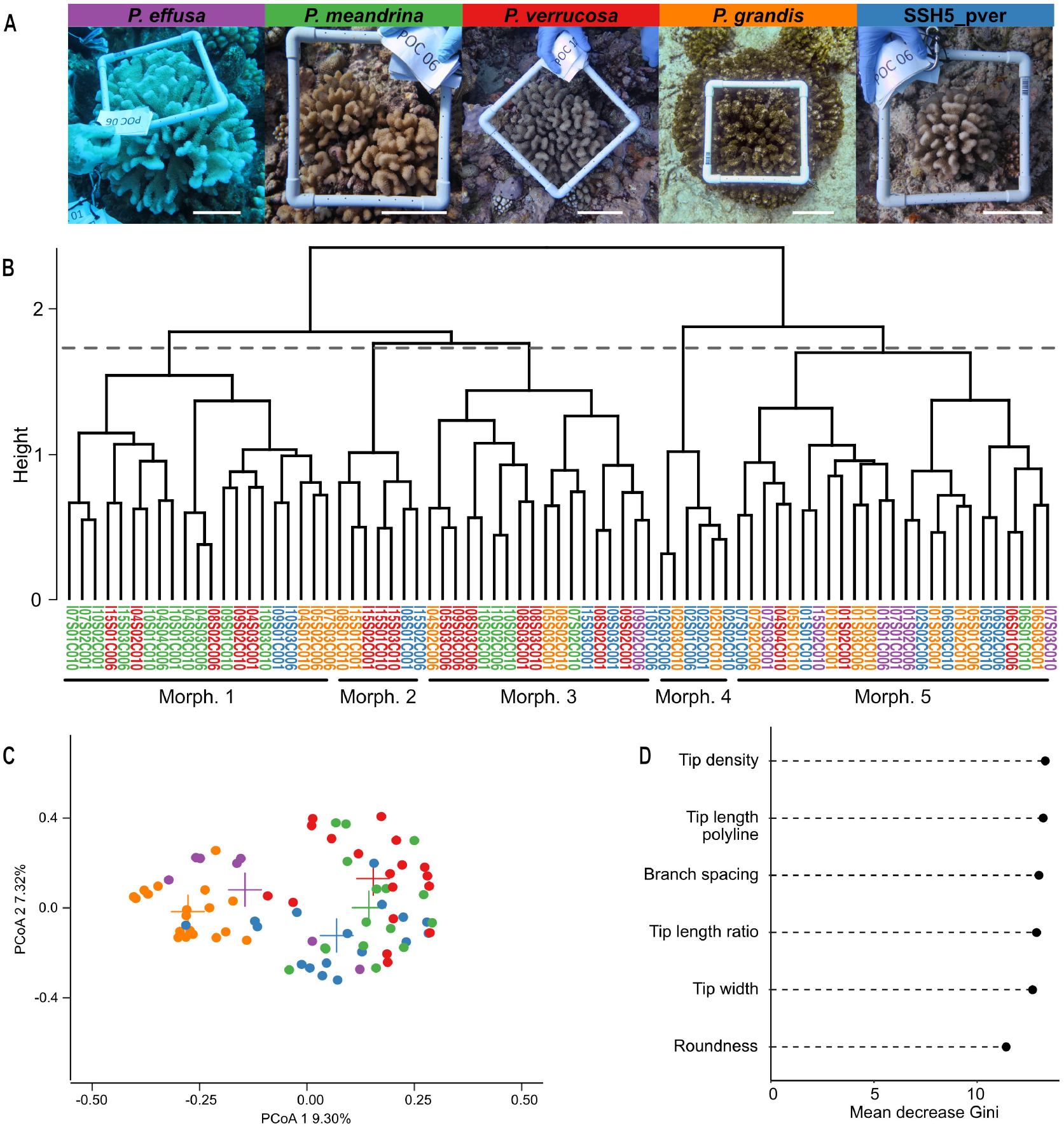
Morphological analysis of *Pocillopora*. A) Representative photos for each of the five SSH. B) Dendrogram of hierarchical clustering based on unsupervised Random Forest analysis. The cutoff used to cluster the samples into five groups (morphotypes) is indicated by the horizontal gray dotted line. Morphotypes are annotated below the dendrogram. C) PCoA generated from the unsupervised Random Forest proximity matrix with samples colored according to the SVD designations given in A and with the centroids of the groups indicated as crosses. D) The mean decrease in Gini test indicating the importance of each variable in determining the homogeneity in the nodes of the decision trees in the unsupervised Random Forest model.

**Supplementary Figure 4.**
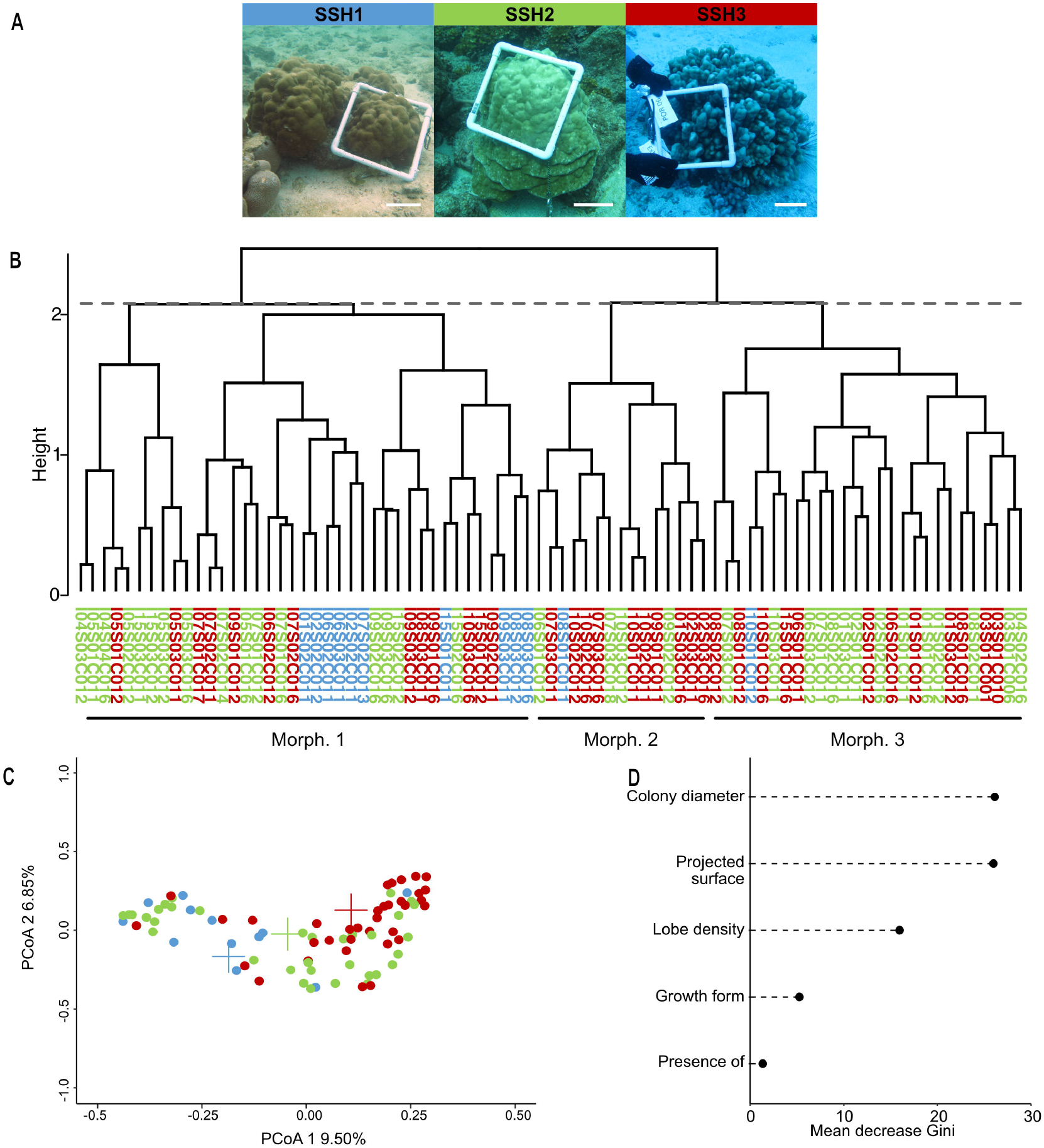
Morphological analysis of *Porites*. A) Representative photos for each of the three SSH. B) Dendrogram of hierarchical clustering based on unsupervised Random Forest analysis. The cutoff used to cluster the samples into three groups (morphotypes) is indicated by the horizontal gray dotted line. Morphotypes are annotated below the dendrogram. C) PCoA generated from the unsupervised Random Forest proximity matrix with samples colored according to the sNMF designations given in A and with the centroids of the groups indicated as crosses. D) The mean decrease in Gini test indicating the importance of each variable in determining the homogeneity in the nodes of the decision trees in the unsupervised Random Forest model.

**Supplementary Figure 5.**
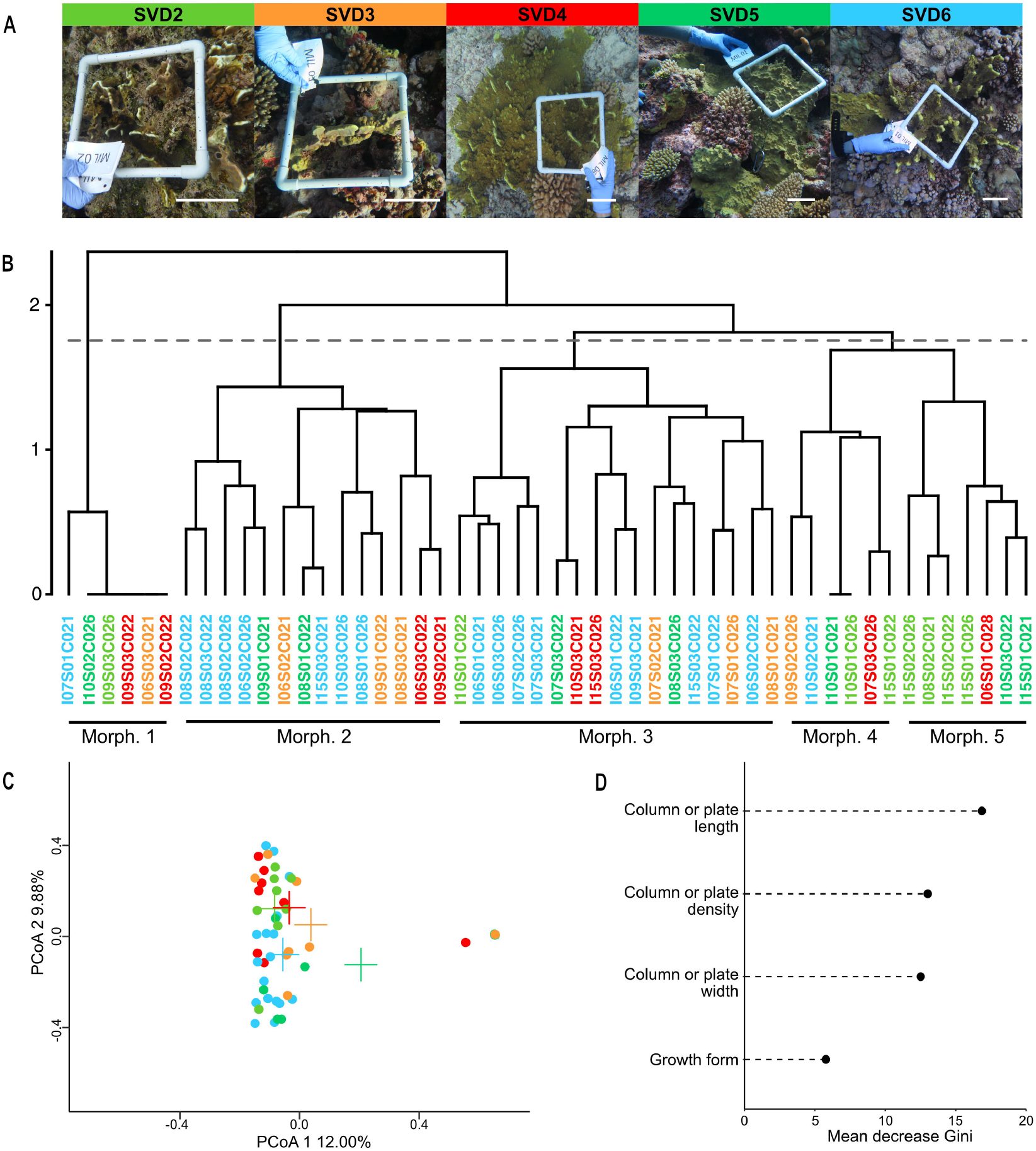
Morphological analysis of *Millepora*. A) Representative photos for each of the five genetic delineations (SVD1 *M. intricata* samples excluded). B) Dendrogram of hierarchical clustering based on unsupervised Random Forest analysis. The cutoff used to cluster the samples into five groups (morphotypes) is indicated by the horizontal gray dotted line. Morphotypes are annotated below the dendrogram. C) PCoA generated from the unsupervised Random Forest proximity matrix with samples colored according to the SVD designations given in A and with the centroids of the groups indicated as crosses. D)The mean decrease in Gini test indicating the importance of each variable in determining the homogeneity in the nodes of the decision trees in the unsupervised Random Forest model.

**Supplementary Figure 6.**
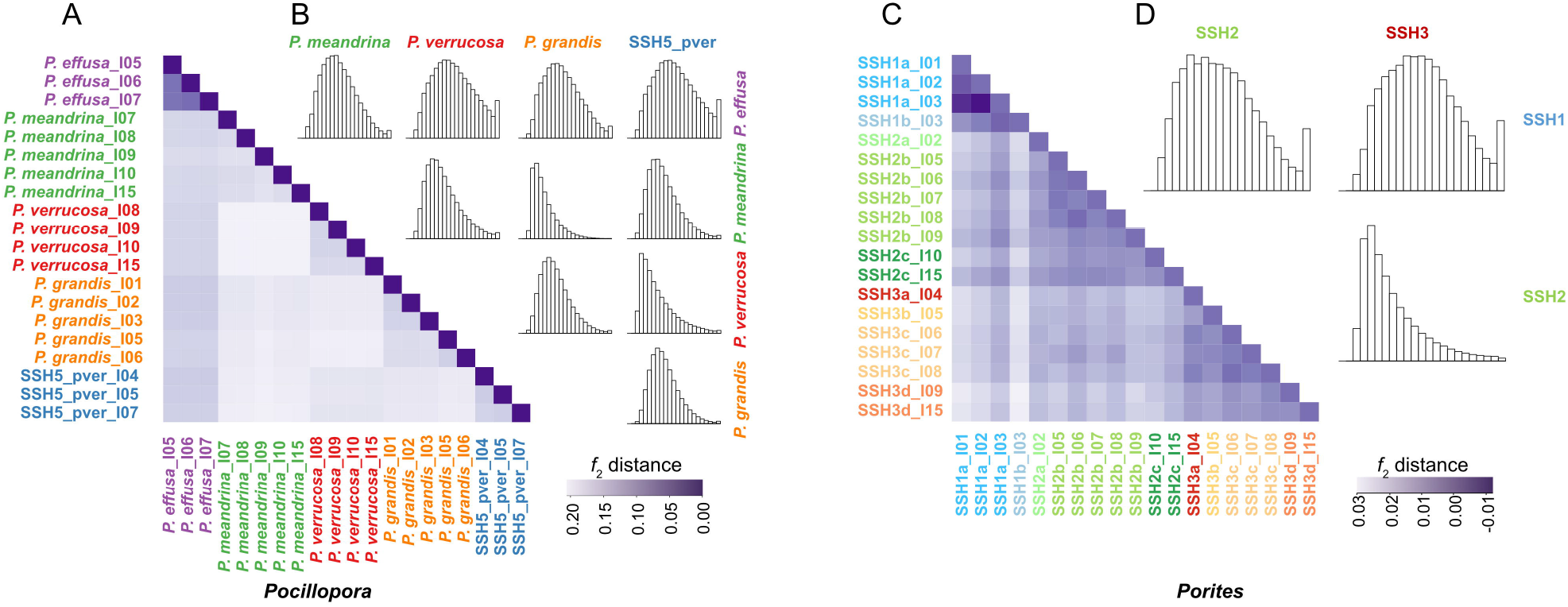
Characterization of among and within SSH genetic differentiation. A & C) *f*_2_ distances indicating correlations in genetic content between *Pocillopora* and *Porites* samples, respectively. B & D) Distributions of between SSH Weir’s *F*_ST_ values calculated in 500 bp sliding windows for *Pocillopora* and *Porites*, respectively. For each histogram the y axis represents abundance while the x axis represents *F*_ST_ value with 0 on the left and 1 on the right.

### SUPPLEMENTARY TABLE LEGENDS

**Supplementary Table 1. Coordinates and number of samples collected at each of 32 sites from 11 islands across the Pacific**.

**Supplementary Table 2. Testing of secondary species hypotheses (SSH) in *Pocillopora* and *Porites* using Bayes factor delimitation with genomic data (BFD*)**. The hypothesis resolving each SVD as a distinct species was the second most likely in each of two replicate runs (Run A and Run B) and is underlined.

**Supplementary Table 3. List of all colonies sampled and their genetic delineations**. The sample names as used in the present study are given in ‘sample name’. ‘TARA barcode’ and ‘sampling-design_label’ provide additional sample identifiers that integrate with the wider *Tara* Pacific dataset. The columns ‘species’ (*Pocillopora*), ‘SSH,subcluster’ (*Porites*), and ‘sNMF,SVDquartet’ (*Millepora*) designate the respective species and genetic delineations. The column ‘MLL (clone)’ denotes membership of the sample to a given clonal Multi Locus Lineage (repeated genet; MLL), and the column “ramets kept” marks if the colony was kept in the clonality pruned dataset (1 ramet per genet). The column BFD* (*Pocillopora* and *Porites* only) marks the replicate BFD* run to which it belongs. The “reference individual” column (*Pocillopora*) marks the samples used for species name attribution. The ‘introgression population’ column gives the population to which the colony was assigned for the introgression analysis.

**Supplementary Table 4. Mantel tests of genetic distances against geographic and historical temperature distances**. Tests are given for each of the genera and for each of the lineages within the genera. Tests for correlation between geographic and historical temperature distances for each of the genera are also given. Significant (p<0.05) results are underlined.

## ONLINE METHODS

### Sample collection, data production, and deposition

Corals were sampled at 11 islands encompassing 32 sites (∼3 sites per island) across an 18,000 km longitudinal transect from July 2016 to January 2017 during the first year of the *Tara* Pacific Expedition (Planes et al. 2019). Coordinates of the sampled sites and the number of samples collected are detailed in Supplementary Table 1. Three target species were sampled based on coral colony morphology: *Pocillopora meandrina, Porites lobata*, and *Millepora* cf. *platyphylla*. In addition, at I02S01 (Island 2, Site 1), 3 colonies of *Millepora intricata (Boissin et al. 2020)*, that displayed a distinct branching morphology, were collected. Where possible, at each site, a minimum of three corals were sampled and photographed as detailed in Lombard et al. 2022 for a total of 106, 109 and 57 colonies of *P. meandrina, P. lobata* and *Millepora* spp., respectively (Supplementary Table 1). At each sampling site environmental data was also collected, and records of historical temperature were associated as detailed in Lombard et al. 2022.

For the *Pocillopora* and *Porites* colonies, total DNA was isolated for each colony fragment as detailed in Belser et al. 2022 and for 3 colonies at each site (totaling 106 and 109 colonies), metagenomic shotgun sequencing was performed with a targeted yield of 1.10^8^ 150 bp paired end reads. For the *Millepora* colonies, due to lack of a reference genome, RNA-Seq was performed as detailed in Belser et al. 2022 on all colonies to enable genotyping using a ‘target gene’ approach (detailed below). Deposition of sequencing data is also detailed in Belser et al. 2022.

### SNP calling and filtering

For *Pocillopora* and *Porites* colonies, we identified a set of genome-wide single nucleotide polymorphisms (SNPs) by mapping the metagenomic reads to the *Pocillopora meandrina* and *Porites lobata* genomic references generated by (Noel et al. 2022) using the Genome Analysis Toolkit program (Van der Auwera and O’Connor 2020) (GATK, v3.7.0). We followed a modified version of the best practices protocol for variant discovery with GATK with manual filtering of the resulting variants. The following protocol was carried out for each species independently.

We aligned Illumina-generated 150-bp paired-end metagenomic reads sequenced from each colony to the predicted coding sequences of the *Pocillopora meandrina* and *Porites lobata* host reference genomes using Burrows–Wheeler Transform Aligner (BWA-mem, v0.7.15) with the default settings (H. Li and Durbin 2009). A read was considered a host contig if its sequence aligned with ≥ 95% sequence identity and with ≥ 50% of the sequence aligned. Host-mapped reads were sorted and filtered to remove sequences which contained > 75% of low-complexity bases and < 30% high-complexity bases using SAMtools v1.10.2 (Heng Li et al. 2009) with the resultant bam files visualized using the Integrated Genomics Viewer (Robinson et al. 2011). The reference genomes were indexed with picardtools’ v2.6.0 (https://broadinstitute.github.io/picard/) *CreateSequenceDictionary* before performing local realignment around small insertions and deletions (indels) using GATK’s RealignerTargetCreator and IndelRealigner to reduce false positive variant identification and represent indels more parsimoniously.

After sample pre-processing, we called DNA variants (SNPs and indels) individually for each colony (GATK, HaplotypeCaller), generating one Genomic Variant Call Format (GVCF) file per sample. We then combined all per-colony GVCF files for a given island (ca. 9 files per island) into a single, multi-sample GVCF file (GATK, *CombineGVCFs*). This resulted in the generation of 11, island-specific combined GVCF files for each species. We then consolidated these 11 multi-sample GVCFs into a GenomicsDB database which allowed for subsequent variant calling across all island cohorts through a joint genotyping analysis (GATK, *GenotypeGVCFs*). Because the GenotypeGVCFs tool is capable of handling any ploidy level (or mix of ploidies) intelligently, we did not specify ploidy level in our function call. This analysis resulted in the generation of a single, “raw” Variant Call Format (VCF) file for each species in which all colonies were jointly genotyped.

Because well-curated filtering resources necessary for GATK’s Variant Quality Score Recalibration (VQSR) tool were not available for either coral species, we filtered the raw VCF files manually using VCFtools (Danecek et al. 2011) v0.1.12 keeping only biallelic sites with minor allele frequencies ≥ 0.05, site-quality scores ≥ 30, and no missing data across colonies. This produced the “linked” datasets (5,937,714 and 1,971,638 variant sites with minimum read depth coverages ≥ 16 and 17 for *Pocillopora* and *Porites*, respectively) from which the SNPs (i.e., discounting indels) were used for the identification of genomic islands of differentiation and of the genomic signature of selection.

In order to avoid variant site linkage effects in downstream genetic analyses, we additionally filtered variants on linkage disequilibrium. We used the ‘prune’ add-on feature in BCFtools (Danecek et al. 2021; H. Li 2011) v1.11 to discard variants with a linkage disequilibrium (r^2^) ≥ 0.2 in a window of 1,000 sites. This resulted in the “unlinked” datasets (347,243 and 183,222 variant sites for *Pocillopora* and *Porites*, respectively). The SNPs from this curated set were used to identify Secondary Species Hypotheses (SSH) using individual based phylogenies, genetic hierarchical clustering and species tree, and to test for population structure, admixture and Genotype Environment associations, as described below.

For the detection of clonal lineages, we had to further reduce the size of these unlinked datasets to avoid unreasonable computing times. This was performed in *Pocillopora* by first further filtering on linkage disequilibrium (r2 ≥ 0.02) by BCFtools to a 27,382 variant site dataset for a first round of clone detection, and then by partitioning the individuals by SVDquartet lineages (see below). The coalescent testing of SSH needed a further size reduction of the datasets, again on linkage disequilibrium and keeping only one individual per putative genetic lineage per island (two replicates of 25 individuals and 636 variant sites, and of 24 individuals and 1,032 variant sites for *Pocillopora* and *Porites*, respectively).

Because no reference genome was available at the time for SNP calling in *Millepora* cf. *platyphylla*, a ‘target gene’ approach was followed. Trimmed RNA-seq metatranscriptomic sequences were subjected to de novo whole transcriptome assembly using Trinity (Grabherr et al. 2011), after being further reduced to 23,755,336 reads by in silico normalization. Trinity assembly produced 302,299 transcripts, which were then clustered into 40,560 unigenes (i.e. uniquely assembled transcripts) with N50 of 1,053 bp. Based on BUSCO, the assembled transcriptome was highly complete with 85.6% of the ortholog genes from the Eukaryota database being present with low fragmentation (3.6%), missing (10.8%), and duplication (16.7%) metrics. Within this transcriptome we identified *Millepora* orthologs for a set of arbitrary stress and/or environment response genes from various cnidarian: stress response genes from scleractinians and actinarians, or genomic fragments from *Pocillopora* and *Porites* that contain temperature outlier SNPs obtained within the present study. The *Millepora* orthologs were obtained using QuickParanoid that takes a collection of files produced by InParanoid (Sonnhammer and Östlund 2015) as input and finds ortholog clusters among multiple species. The protocol used strictly followed the instructions published in http://pl.postech.ac.kr/QuickParanoid/. Briefly, for a given set of “target genes”, QuickParanoid first preprocesses each InParanoid output file by using Blastall and then computes ortholog clusters. As for the previous coral genera, metagenomic reads were aligned on this set of target genes using Burrows–Wheeler Transform Aligner (BWA-mem, v0.7.15) with the default settings (Li and Durbin 2009). Host-mapped reads were then sorted and processed using SAMtools v1.10.2 (Li et al. 2009) to generate respective bam files and SNPs were identified as previously with GATK. The list of orthologs for the *Millepora* transcriptome contigs that contained SNPs is part of the resource files and scripts deposited in (Hume et al. 2022). The SNP dataset was then filtered for biallelic status, SNPs only, quality, missingness and minimum allele frequency with the same thresholds as for the other corals resulting in a final set of 446 variant sites. One individual with an excess of missing data (I09S03C026) was not included in the sNMF analyses. Filtering on linkage disequilibrium (r^2^>0.2) for further genetic analyses reduced this dataset to 243 ‘unlinked” variant sites in 56 individuals.

### Species delimitation overview

Here we followed the integrative taxonomy approach outlined by Pante *et al*., 2015 (Eric Pante et al. 2015) by treating our morphology-guided sampling of the three coral species as primary species hypotheses (PSH) from which we developed secondary species hypotheses (SSH) through analysis of genetic diversity and colony macromorphology. We then formally tested the SSH using coalescent analysis (Carstens et al. 2013).

### Genetic delineation of species

To get an initial impression of the genetic diversity contained in the *Pocillopora* and *Porites* samples we built an individual-based Maximum Likelihood phylogenetic tree using RAxML (Stamatakis 2014). For this, the “unlinked SNPs” vcfs were translated to relaxed Phylip format using PGDSpider (Lischer and Excoffier 2012), and raxmlHPC-PTHREADS was run on 40 cores, with a GTRCAT model and the number of bootstrap replicates determined by the autoFC option. The best ML tree with support value was then transferred in Newick format to MEGA X (Kumar et al. 2018) for graphical annotation.

A species tree was then built using SVDquartets (Chifman and Kubatko 2014) as implemented in PAUP* (Rédei 2008) v4.0a152 for all three genera. This nonBayesian approach infers the relationship among quartets of taxa under a coalescent model and uses this information to build the species tree. Parallel computing on 50 threads was performed to analyze all possible quartets and 100 bootstrap replicates. The bootstrap consensus tree was then transferred to Mega X for graphical annotation.

To select which of the SVDquartet clades formed pertinent SSH, a hierarchical genetic clustering was also performed using the program sNMF (Frichot et al. 2014) with the snmf function from the LEA (Frichot and François 2015) R package. However, as this analysis is sensitive to clonality (B. Porro pers. comm.), we first identified clonal lineages in the *Pocillopora, Porites* and *Millepora* “unlinked” datasets using the R package Rclone (Bailleul, Stoeckel, and Arnaud⁏Haond 2016) to identify the pairwise threshold distance between clonal individuals, and used mlg.filter function from the R package poppr (Kamvar, Tabima, and Grünwald 2014; Kamvar, Brooks, and Grünwald 2015) to assign individuals to clonal, or, more properly, multilocus lineages (MLL). We ran this clonal analysis on all samples for each of the 3 genera. In *Pocillopora* and *Porites*, due to a considerable lineage-correlating variation in pairwise distances, we also ran further analyses on SVD-grouping-defined subsets of samples. The subsequent sNMF analyses were performed keeping only one ramet per clonal genet with the exception of the 3 *M. intricata* samples that were identified as being a single MLL. sNMF determines the optimal number of ancestral populations from which the actual dataset could be issued through admixture. These analyses were performed on 50 cores and 10 repetitions for a number of possible ancestral populations that varied from 1 to 20, with an alpha parameter value of 10. The best number of ancestral clusters was determined by the entropy criterion and a bar chart representing the individual admixture coefficient from each of these ancestral clusters was produced. To test for possible sub clustering, a second round of snmf analysis was performed separately on each of the ancestral clusters identified in the first round (with the same analysis parameters). As a further visualization of the genetic diversity within each of the 3 genera, we performed principal component analysis (PCA) based on individual genome wide genotypes using the glPca function of the adegenet R package (Jombart and Ahmed 2011; Jombart 2008). For *Pocillopra* and *Porites*, sNMF clustering and SVDquartet species tree topology then guided the construction of secondary species hypotheses to be tested by coalescent analysis using Bayes factor delimitation with genomic data (BFD*) (Grummer, Bryson, and Reeder 2014; Bryant et al. 2012). BFD analyses were not conducted in *Millepora* due to the low number of SNPs. The BFD* analyses were performed in BEAST2 (Bouckaert et al. 2019) with 48 path sampler sets of 500,000 MCMC repetitions, to sample in 10,000,000 MCMC iterations of SNAPP (Bryant et al. 2012) for *Pocillopora* and 750,000 MCMC repetitions and 1,000,000 MCMC iterations of SNAPP for *Porites*. These numbers of iterations were determined as minimal for convergence by first running standard SNAPP analyses in Beast2. To select for the most pertinent SSH, the BFD* likelihood ranking of the SSH was performed twice in each genus on a duplicate set of individuals for the same variant sites.

### Morphology analysis

To investigate the degree to which morphology may support or confound the sofar genetics-based SSH we categorized the samples into morphotypes through morphological analysis and contrasted these groupings with the genetic resolutions. The *M. intricata* samples were excluded from the analysis. Photographs of entire colonies were taken for all samples using an underwater camera with 20 × 20 cm quadrats as a scale (photographs available online at https://store.pangaea.de/Projects/TARA-PACIFIC/Images/). Collection of morphological parameters was conducted using Imagej (Schindelin et al. 2012), the image annotation plugin objectj (https://sils.fnwi.uva.nl/bcb/objectj/), and customs macros with the photos corrected for perspective using the “interactive perspective” plugin with the quadrat as a reference.

The following morphological parameters were collected: For *Pocillopora*: branch tips density (calculated from the number of tips divided by the projected surface of the whole colony), tip length polyline (a line connecting the two extremities of the top of a branch following the curvature of the tip; n = 3), tip length ratio (the ratio between a straight line connecting the extremities of the top of a branch and the tip length polyline to detect the degree of meander shape of the branch tip; n = 3), tip width (n = 3), and colony roundness (the difference between the maximum and minimum diameters, and branch spacing between a randomly selected center branch and six adjacent branches; n = 6). For *Porites*: colony diameter, projected surface, growth form (columnar, massive or encrusting), lobe density (computed by the number lobes divided by the area of a 20 × 20 cm quadrat drawn on the colony photo using the Imagej rectangle tool), and presence of ridges (present or absent). For *Millepora*: growth form (encrusting, laminar, columnar, branching columnar), column or plate density (computed by the number of columns or plates within the 20 × 20 cm quadrat), length of columns or plates (the length of a straight line connecting the extremities of the top of a column/plate; n = 1 - 6, when present), and width of columns/plates (n = 1-6, when present).

Morphometric analysis of the morphological parameters was conducted in R V4.1.0 using the same approach for all three genera. Unsupervised Random Forest analysis (randomForest::randomforest) was used to generate a proximity matrix upon which hierarchical clustering was performed to identify morphotypes (stat::hclust, factoextra::fviz_dend) with a threshold set so that the number of clusters equated to the number of genetically derived SSH (*Pocillopora* and *Porites*), or the six SVD groupings (*Millepora*). The Gini coefficient was tested to measure the contribution of morphological variables in the homogeneity of the nodes of the decision trees. The proximity matrix was used as input to generate a principal coordinate analysis ordination (PCoA; stats::cmdscale) to visualize variance in morphology across the samples in accordance with the genetic lineage assignments. Additionally, PERMANOVA (pairwiseAdonis::pairwise.adonis2; 999 permutations; p-adjusted Bonferroni correction applied; https://github.com/pmartinezarbizu/pairwiseAdonis) was used to test whether variance in morphology could be significantly explained by the genetic delineations.

### Identification of *Pocillopora* SSH taxonomy

To assign species names to our SSH, two representatives for each SVD were mapped on the 2,068 *Pocillopora* reference sequences from (Oury et al. 2022) using BWA (H. Li and Durbin 2009). Samples were then genotyped for the 1,559 unlinked diagnostic SNPs generated therein. Genotypes with a minimum read depth (DP) of 12× and non-significant strand biases (SP < 13) were called and filtered with BCFtools (H. Li 2011; Danecek et al. 2021) v1.11. Between 1,489 and 1,524 SNPs were retrieved per sample. The resulting vcf was then combined with the one used for species delimitation in (Oury et al. 2022), and assignment tests were performed from *K* = 2 to *K* = 30 using the program sNMF (Frichot et al. 2014) with the snmf function from the LEA (Frichot and François 2015) R package, as previously described. From that, SSH correspondance with genomic species hypotheses (GSH) from (Oury et al. 2022), and thus current taxonomy, was retrieved. Such resources were not available for *Porites* and insufficient SNPs limited such an analysis for *Millepora*.

### Among and within species genetic differentiation

To further resolve evolutionary history in the *Pocillopora* and *Porites* species designations, we investigated introgression and pairwise divergence for the SSH. Following Louis *et al*. 2021 (Louis et al. 2021), introgression among lineages was computed with TreeMix (Pickrell and Pritchard 2012). The unlinked SNP dataset vcf was converted to TreeMix format using vcftools (Danecek et al. 2011) and PLINK (Purcell et al. 2007). The individuals were regrouped in populations according to their SSH and islands of origin, and the optimal number of admixture events was estimated at this level. For this we ran TreeMix 10 times for 0 to 10 events and estimated the optimal number of events using the optM R packages (Fitak 2021). We then ran TreeMix 100 times for 0 to this optimal number of admixture/introgression events to produce a consensus tree and bootstrap values using the BITE R package (Milanesi et al. 2017). The residual covariance matrix was estimated for the optimal number of admixture/introgression events and the consensus tree using TreeMix. The occurrence of the admixture/introgression events were verified by computing the relevant *f*_4_ indices (Reich et al. 2009) and correlation in genetic content was further quantified by calculating pairwise *f*_2_ distances both using the AdmixTools2 R package (https://uqrmaie1.github.io/admixtools/index.html) after converting the unlinked SNP dataset vcf to the BED format using vcftools (Danecek et al. 2011) and PLINK (Purcell et al. 2007).

To quantify and characterize genomic divergence between the SSH within each of *Pocillopora* and *Porites* we computed pairwise Weir and Cockerham (Weir and Cockerham 1984) *F*_ST_ along the genome (500 bp sliding window) with vcftools (Danecek et al. 2011).

### Correlation of genetic lineages with geographic distance and historical thermal regime

To assess for isolation-by-distance and a possible effect of temperature regime on the evolutionary trajectories of the species, we conducted correlation analyses sensu Rousset 1997 (Rousset 1997). For *Pocillopora* and *Porites*, samples were again grouped according to their SSH and islands of origin (i.e. SVD4_I01, SVD4_I02… etc.) while *Millepora* samples were grouped according to island of orgin. Within each genus, pairwise group *F*_ST_ distances were generated using vcftools (Danecek et al. 2011) and used to generate pairwise matrices of *F*_ST_/(1-*F*_ST_). These distances, representative of genetic dissimilarity, were then tested for correlation against distance matrices representing geographic and historical temperature (environmental) distances using the mantel function from the vegan R package (Oksanen et al. 2020). Geographic distances were based on GPS coordinates of islands (estimated by an average of the site coordinates). Euclidean historical temperature distances were generated from a multiparameter historical temperature dataset collected for each site as part of the *Tara* Pacific consortium (as detailed in Lombard et al. 2022) (Lombard et al. 2022) and used in a site-wise manner after conducting PCA to reduce its dimensionality from 63 initial parameters to the 5 highest scoring PCs that cumulatively accounted for >80% of the observed variance (using R package FactoMineR) (Lê, Josse, and Husson 2008).

### Effect of environment on SSH divergence

To assess the influence of environment (using historical temperature as a proxy) on divergence between the SSH in *Pocillopora* and *Porites* we first used the reduced dimension historical temperature dataset (see above) to identify temperature outlier SNPs whose allelic frequencies were linked to historical temperature variation using redundancy analysis (RDA) following the exact protocol of Capblancq and Forester (Capblancq and Forester 2021). For this, the unlinked SNP dataset had first to be converted to ped format using vcftools. From this RDA, we identified temperature outlier SNPs, *i*.*e*. the SNPs whose allelic frequencies distribution among populations were best explained by the local historical temperature variation, as the 100 SNPs with the associated lowest *p*-values (respectively *p* < 2.15 10^−4^ in *Pocillopora* and *p* < 1.72 10^−3^ in *Porites*). We then defined a set of genomic islands of differentiation (GID) as the 1% of the 500bp bins with the highest mean *F*_ST_ value after conducting a further pairwise sliding window *F*_*ST*_ analysis as detailed above but with samples grouped according to SSH for each of the genera. We compared the proportion of these temperature outlier SNPs that were found in GIDs (see above) between the two genera. Finally, we investigated the proportion of the temperature outlier SNPs, and in particular those located in GIDs, that were predicted to be under divergent selection using the program Flink (Galimberti et al. 2020). Flink operates on a dataset of linked loci and is based on the F-model which measures differences in allele frequencies (*F*_ST_) that are partitioned into locus- and population-specific components reflecting selection and drift, respectively. Flink structures samples in a hierarchical model with the level of ‘population’ nested in ‘group’ and ‘group’ nested in ‘higher hierarchy’. We equated ‘group’ to the SSH, and ‘population’ to island of collection for the samples and generated allele counts from the ‘linked SNPs’ dataset as input using ATLAS (Link et al. 2017). To reduce computational complexity, the target genome can be subdivided with individual Flink analysis run across each subdivision (Galimberti et al. 2020). However, each subdivision must have a suitable number of variant sites to enable effective inference of demographic parameters with 10,000 suggested as a minimum by the authors. As such we first ran one instance of Flink for every contig in the *Pocillopora* and *Porites* genomes with at least 10,000 variants (subsampling using BCFtools (Danecek et al. 2021) v1.9) using 4 cores per instance and a total of 800,000 MCMC iterations (300,000 burnin). Analysis showed that the demographic parameters converged at similar values across contigs within each of the respective genomes. To identify loci under selection in the remaining contigs (i.e. those with < 10,000 variants) we therefore ran one instance of Flink per contig with parameters fixed according to the converged demographic parameters of the previous runs. Contig-wise results were consolidated and loci under divergent selection at the higher hierarchy level (i.e. between lineages; output file ‘Posterior_A.txt’) were assessed by calculating a false discovery rate (FDR) from the posterior alpha distributions following the tool’s documentation using a custom python script. The Nextflow (Di Tommaso et al. 2017) pipelines used to run the analyses and the Python and R scripts used to process the results are available on GitHub (https://github.com/didillysquat/TARA_host_popgen_2022).

## Data and Script availability

A collection of resource files and scripts used to conduct the analyses, including the raw and unfiltered vcf files, have been deposited as a Zenodo deposition (https://doi.org/10.5281/zenodo.7229405; Hume et al. 2022).

