## Supplementary_Table_1 for "Disparate patterns of genetic divergence in three widespread corals across a pan-Pacific environmental gradient highlights species-specific adaptation trajectories"

**Supplementary Table 1.** **Coordinates and number of samples collected at each of 32 sites from 11 islands across the Pacific.**

| **Island number** | **Island name** | **Site number** | **Latitude** | **Longitude** | ***Pocillopora*** | ***Porites*** | ***Millepora*** |
| --- | --- | --- | --- | --- | --- | --- | --- |
| 1 | Las Perlas | 1 | 8.5795 | -79.02055 | 3 | 3 | 0 |
| 1 | Las Perlas | 2 | 8.597875 | -79.024997 | 5 | 3 | 0 |
| 1 | Las Perlas | 3 | 8.6501 | -79.0339 | 3 | 3 | 0 |
| 2 | Coiba | 1 | 7.8732 | -81.7942 | 4 | 3 | 3 |
| 2 | Coiba | 2 | 7.6513 | -81.697 | 3 | 3 | 0 |
| 2 | Coiba | 3 | 7.2092 | -81.7962 | 3 | 3 | 0 |
| 3 | Malpelo | 1 | 3.9873 | -81.5915 | 10 | 12 | 0 |
| 4 | Easter | 1 | -27.0688 | -109.3233 | 3 | 3 | 0 |
| 4 | Easter | 2 | -27.0674 | -109.335 | 3 | 3 | 0 |
| 4 | Easter | 3 | -27.1326 | -109.434 | 3 | 3 | 0 |
| 4 | Easter | 4 | -27.1486 | -109.4442 | 3 | 3 | 0 |
| 5 | Ducie | 1 | -24.6971 | -124.8021 | 3 | 3 | 0 |
| 5 | Ducie | 2 | -24.6971 | -124.7947 | 3 | 3 | 0 |
| 5 | Ducie | 3 | -24.6709 | -124.7757 | 3 | 3 | 0 |
| 6 | Gambier | 1 | -23.0748 | -135.072 | 3 | 1 | 3 |
| 6 | Gambier | 2 | -23.1651 | -134.8482 | 3 | 3 | 3 |
| 6 | Gambier | 3 | -23.2374 | -134.9566 | 3 | 3 | 3 |
| 7 | Moorea | 1 | -17.482567 | -149.886467 | 3 | 4 | 3 |
| 7 | Moorea | 2 | -17.518433 | -149.924 | 3 | 5 | 3 |
| 7 | Moorea | 3 | -17.489667 | -149.75505 | 3 | 4 | 3 |
| 8 | Aitutaki | 1 | -18.839967 | -159.8009 | 3 | 3 | 3 |
| 8 | Aitutaki | 2 | -18.913967 | -159.8451 | 3 | 3 | 3 |
| 8 | Aitutaki | 3 | -18.8678 | -159.818667 | 3 | 3 | 3 |
| 9 | Niue | 1 | -19.113433 | -169.914233 | 3 | 3 | 3 |
| 9 | Niue | 2 | -18.985417 | -169.903267 | 3 | 3 | 3 |
| 9 | Niue | 3 | -19.041817 | -169.9185 | 3 | 3 | 3 |
| 10 | Samoa | 1 | -14.010967 | -171.843117 | 3 | 3 | 3 |
| 10 | Samoa | 2 | -14.0616 | -171.430367 | 3 | 3 | 3 |
| 10 | Samoa | 3 | -13.918767 | -171.541567 | 3 | 3 | 3 |
| 15 | Guam | 1 | 13.249733 | 144.64495 | 3 | 3 | 3 |
| 15 | Guam | 2 | 13.4172 | 144.644633 | 3 | 3 | 3 |
| 15 | Guam | 3 | 13.3434 | 144.6361 | 3 | 3 | 3 |
| Totals |  |  |  |  | 106 | 109 | 57 |
