## Supplementary_Table_2 for "Disparate patterns of genetic divergence in three widespread corals across a pan-Pacific environmental gradient highlights species-specific adaptation trajectories"

**Supplementary Table 2*.* *Pocillopora* and *Porites* BFD***: Testing of secondary species hypotheses (SSH) using Bayes factor delimitation with genomic data (BFD*). The hypothesis resolving each SVD as a distinct species was the second most likely in each of two replicate runs (Run A and Run B) for each species and is underlined.

| **Run A** | | **Run B** | |
| --- | --- | --- | --- |
| **Species split (nb. species)** | **Marginal Likelihood** | **Species split (nb. species)** | **Marginal Likelihood** |
| ***Pocillopora*** |  |  |  |
| SVD1/SVD2/SVD4/SVD3-SVD5 (4) | -12963.059 | SVD1/SVD2-SVD4/SVD3-SVD5 (3) | -12883.316 |
| SVD1/SVD2/SVD3/SVD4/SVD5 (5) | -12973.734 | SVD1/SVD2/SVD3/SVD4/SVD5 (5) | -12895.469 |
| SVD1/SVD2-SVD4/SVD3-SVD5 (3) | -12979.111 | SVD1/SVD2 /SVD4/SVD3-SVD5 (4) | -12902.967 |
| SVD1/SVD2-SVD4/SVD3/SVD5 (4) | -13005.359 | SVD1/SVD2-SVD4/SVD3/SVD5 (4) | -13001.632 |
| SVD1/SVD2-SVD3-SVD4-SVD5 (2) | -13193.471 | SVD1/SVD2-SVD3-SVD4-SVD5 (2) | -13075.553 |
| ***Porites*** |  |  |  |
| K1a/K1b/K2abc/K3abcd (4) | -9149.427 | K1ab/K2a/K2b/K2c/K3abcd (5) | -9181.521 |
| K1ab/K2abc/K3abcd (3) | -9154.839 | K1ab/K2abc/K3abcd (3) | -9188.583 |
| K1ab/K2a/K2b/K2c/K3abcd (5) | -9174.805 | K1a/K1b/K2a/K2bc/K3a/K3bcd (6) | -9193.623 |
| K1a/K1b/K2a/K2bc/K3a/K3bcd (6) | -9178.793 | K1a/K1b/K2abc/K3abcd (4) | -9193.789 |
| K1ab/K2abc/K3a/K3b/K3c/K3d (6) | -9191.031 | K1ab/K2abc/K3a/K3b/K3c/K3d (6) | -9210.469 |
