## Supplementary_Table_3 for "Disparate patterns of genetic divergence in three widespread corals across a pan-Pacific environmental gradient highlights species-specific adaptation trajectories"

**Supplementary Table 3: List of all colonies sampled and their genetic delineations.** The sample names as used in the present study are given in ‘sample name’. ‘TARA barcode’ and ‘sampling-design_label’ provide additional sample identifiers that integrate with the wider TARA Pacific dataset. The columns ‘species (*Pocillopora*), ‘SSH,subcluster’ (*Porites*), and ‘sNMF,SVDquartet’ (*Millepora*) designate the respective species and genetic delineations. The column ‘MLL (clone)’ denotes membership of the sample to a given clonal Multi Locus Lineage (repeated genet; MLL), and the column “ramets kept” marks if the colony was kept in the clonality pruned dataset (1 ramet per genet). The column BFD* (*Pocillopora* and *Porites* only) marks the replicate BFD* run to which it belongs. The “reference individual” column (*Pocillopora*) marks the samples used for species name attribution*.* The ‘introgression population’ column gives the population to which the colony was assigned to for the introgression analysis.

| **sample name** | **TARA barcode** | **sampling-design_label** | **species** | **introgression population** | **MLL (clone)** | **ramets kept** | **BFD*** | **reference individual** |
| --- | --- | --- | --- | --- | --- | --- | --- | --- |
| ***Pocillopora*** |  |  |  |  |  |  |  |  |
| I01S01C001POC | TARA_CO-0000002 | OA000-I01-S01-C001 | *P. grandis* | SVD4_I01 |  | x |  |  |
| I01S01C006POC | TARA_CO-0000019 | OA000-I01-S01-C006 | *P. grandis* | SVD4_I01 | 1_10 |  | A |  |
| I01S01C010POC | TARA_CO-0000032 | OA000-I01-S01-C010 | *P. grandis* | SVD4_I01 | 1_10 | x | B | x |
| I01S02C001POC | TARA_CO-0000065 | OA000-I01-S01-C001 | *P. grandis* | SVD4_I01 |  | x |  |  |
| I01S02C006POC | TARA_CO-0000070 | OA000-I01-S02-C006 | *P. grandis* | SVD4_I01 |  | x |  |  |
| I01S02C010POC | TARA_CO-0000074 | OA000-I01-S02-C010 | *P. grandis* | SVD4_I01 |  | x | A |  |
| I01S02C021POC | TARA_CO-0000125 | OA000-I01-S02-C021 | *P. grandis* | SVD4_I01 | 1_28 | x |  |  |
| I01S02C022POC | TARA_CO-0000126 | OA000-I01-S02-C022 | *P. grandis* | SVD4_I01 | 1_28 |  |  |  |
| I01S03C001POC | TARA_CO-0000495 | OA000-I01-S03-C001 | *P. grandis* | SVD4_I01 |  | x | B |  |
| I01S03C006POC | TARA_CO-0000500 | OA000-I01-S03-C006 | *P. grandis* | SVD4_I01 |  | x |  |  |
| I01S03C010POC | TARA_CO-0000504 | OA000-I01-S03-C010 | *P. grandis* | SVD4_I01 |  | x |  |  |
| I02S01C001POC | TARA_CO-0000140 | OA000-I02-S01-C001 | *P. grandis* | SVD4_I02 |  | x |  |  |
| I02S01C006POC | TARA_CO-0000145 | OA000-I02-S01-C006 | *P. grandis* | SVD4_I02 |  | x | A+B |  |
| I02S01C010POC | TARA_CO-0000149 | OA000-I02-S01-C010 | *P. grandis* | SVD4_I02 |  | x |  |  |
| I02S01C026POC | TARA_CO-0000205 | OA000-I02-S01-C026 | *P. grandis* | SVD4_I02 |  |  |  |  |
| I02S02C001POC | TARA_CO-0000220 | OA000-I02-S02-C001 | *P. grandis* | SVD4_I02 |  | x |  |  |
| I02S02C006POC | TARA_CO-0000225 | OA000-I02-S02-C006 | *P. grandis* | SVD4_I02 |  | x | B |  |
| I02S02C010POC | TARA_CO-0000229 | OA000-I02-S02-C010 | *P. grandis* | SVD4_I02 |  | x |  |  |
| I02S03C001POC | TARA_CO-0000385 | OA000-I02-S03-C001 | *P. grandis* | SVD4_I02 |  | x |  |  |
| I02S03C006POC | TARA_CO-0000390 | OA000-I02-S03-C006 | *P. grandis* | SVD4_I02 |  | x |  |  |
| I02S03C010POC | TARA_CO-0000394 | OA000-I02-S03-C010 | *P. grandis* | SVD4_I02 |  | x | A |  |
| I03S01C011POC | TARA_CO-0000555 | OA000-I03-S01-C011 | *P. grandis* | SVD4_I03 | 1_26 |  | A |  |
| I03S01C012POC | TARA_CO-0000556 | OA000-I03-S01-C012 | *P. grandis* | SVD4_I03 | 1_26 | x | B |  |
| I03S01C013POC | TARA_CO-0000557 | OA000-I03-S01-C013 | *P. grandis* | SVD4_I03 |  | x |  |  |
| I03S01C014POC | TARA_CO-0000558 | OA000-I03-S01-C014 | *P. grandis* | SVD4_I03 |  | x |  |  |
| I03S01C015POC | TARA_CO-0000559 | OA000-I03-S01-C015 | *P. grandis* | SVD4_I03 | 1_24 | x |  |  |
| I03S01C016POC | TARA_CO-0000560 | OA000-I03-S01-C016 | *P. grandis* | SVD4_I03 | 1_24 |  |  |  |
| I03S01C017POC | TARA_CO-0000561 | OA000-I03-S01-C017 | *P. grandis* | SVD4_I03 | 1_8 | x | A |  |
| I03S01C018POC | TARA_CO-0000562 | OA000-I03-S01-C018 | *P. grandis* | SVD4_I03 | 1_8 |  |  |  |
| I03S01C019POC | TARA_CO-0000563 | OA000-I03-S01-C019 | *P. grandis* | SVD4_I03 | 1_8 |  | B |  |
| I03S01C020POC | TARA_CO-0000564 | OA000-I03-S01-C020 | *P. grandis* | SVD4_I03 | 1_8 |  |  |  |
| I04S01C001POC | TARA_CO-0000580 | OA000-I04-S01-C001 | SSH5_pver | SVD5_I04 |  | x |  |  |
| I04S01C006POC | TARA_CO-0000585 | OA000-I04-S01-C006 | SSH5_pver | SVD5_I04 |  | x |  |  |
| I04S01C010POC | TARA_CO-0000589 | OA000-I04-S01-C010 | SSH5_pver | SVD5_I04 |  | x | B |  |
| I04S02C001POC | TARA_CO-0000655 | OA000-I04-S02-C001 | SSH5_pver | SVD5_I04 |  | x | A |  |
| I04S02C006POC | TARA_CO-0000660 | OA000-I04-S02-C006 | SSH5_pver | SVD5_I04 |  | x |  |  |
| I04S02C010POC | TARA_CO-0000664 | OA000-I04-S02-C010 | SSH5_pver | SVD5_I04 |  | x |  |  |
| I04S03C001POC | TARA_CO-0000697 | OA000-I04-S03-C001 | SSH5_pver | SVD5_I04 |  | x | A |  |
| I04S03C006POC | TARA_CO-0000702 | OA000-I04-S03-C006 | SSH5_pver | SVD5_I04 |  | x |  |  |
| I04S03C010POC | TARA_CO-0000706 | OA000-I04-S03-C010 | SSH5_pver | SVD5_I04 |  | x |  |  |
| I04S04C001POC | TARA_CO-0000777 | OA000-I04-S04-C001 | SSH5_pver | SVD5_I04 |  | x | B |  |
| I04S04C006POC | TARA_CO-0000782 | OA000-I04-S04-C006 | SSH5_pver | SVD5_I04 |  | x |  |  |
| I04S04C010POC | TARA_CO-0000786 | OA000-I04-S04-C010 | SSH5_pver | SVD5_I04 |  | x |  |  |
| I05S01C001POC | TARA_CO-0000817 | OA000-I05-S01-C001 | *P. grandis* | SVD4_I05 |  | x | B |  |
| I05S01C006POC | TARA_CO-0000822 | OA000-I05-S01-C006 | *P. grandis* | SVD4_I05 |  | x |  |  |
| I05S01C010POC | TARA_CO-0000826 | OA000-I05-S01-C010 | *P. effusa* | SVD1_I05 |  | x | B |  |
| I05S02C002POC | TARA_CO-0000919 | OA000-I05-S02-C002 | *P. effusa* | SVD1_I05 |  | x |  |  |
| I05S02C006POC | TARA_CO-0000923 | OA000-I05-S02-C006 | *P. effusa* | SVD1_I05 | 0_5 | x | A | x |
| I05S02C010POC | TARA_CO-0000927 | OA000-I05-S02-C010 | *P. effusa* | SVD1_I05 | 0_5 |  |  |  |
| I05S03C001POC | TARA_CO-0000948 | OA000-I05-S03-C001 | SSH5_pver | SVD5_I05 |  | x | B | x |
| I05S03C006POC | TARA_CO-0000953 | OA000-I05-S03-C006 | SSH5_pver | SVD5_I05 |  | x | A |  |
| I05S03C010POC | TARA_CO-0000957 | OA000-I05-S03-C010 | *P. grandis* | SVD4_I05 |  | x | A | x |
| I06S01C001POC | TARA_CO-0002056 | OA000-I06-S01-C001 | *P. effusa* | SVD1_I06 |  | x | B |  |
| I06S01C006POC | TARA_CO-0002061 | OA000-I06-S01-C006 | *P. effusa* | SVD1_I06 |  | x |  |  |
| I06S01C010POC | TARA_CO-0002065 | OA000-I06-S01-C010 | *P. grandis* | SVD4_I06 |  | x | B |  |
| I06S02C001POC | TARA_CO-0002126 | OA000-I06-S02-C001 | *P. effusa* | SVD1_I06 |  | x |  |  |
| I06S02C006POC | TARA_CO-0002131 | OA000-I06-S02-C006 | *P. effusa* | SVD1_I06 |  | x |  |  |
| I06S02C010POC | TARA_CO-0002135 | OA000-I06-S02-C010 | *P. grandis* | SVD4_I06 |  | x | A |  |
| I06S03C001POC | TARA_CO-0002286 | OA000-I06-S03-C001 | *P. effusa* | SVD1_I06 |  | x |  |  |
| I06S03C006POC | TARA_CO-0002291 | OA000-I06-S03-C006 | *P. effusa* | SVD1_I06 |  | x |  |  |
| I06S03C010POC | TARA_CO-0002295 | OA000-I06-S03-C010 | *P. effusa* | SVD1_I06 |  | x | A |  |
| I07S01C001POC | TARA_CO-0002356 | OA000-I07-S01-C001 | SSH5_pver | SVD5_I07 |  | x |  |  |
| I07S01C006POC | TARA_CO-0002361 | OA000-I07-S01-C006 | SSH5_pver | SVD5_I07 |  | x |  |  |
| I07S01C010POC | TARA_CO-0002365 | OA000-I07-S01-C010 | SSH5_pver | SVD5_I07 |  | x | B |  |
| I07S02C001POC | TARA_CO-0002456 | OA000-I07-S02-C001 | SSH5_pver | SVD5_I07 |  | x |  |  |
| I07S02C006POC | TARA_CO-0002461 | OA000-I07-S02-C006 | SSH5_pver | SVD5_I07 |  | x | A |  |
| I07S02C010POC | TARA_CO-0002465 | OA000-I07-S02-C010 | *P. meandrina* | SVD2_I07 |  | x | A+B |  |
| I07S03C001POC | TARA_CO-0002566 | OA000-I07-S03-C001 | SSH5_pver | SVD5_I07 |  | x |  | x |
| I07S03C006POC | TARA_CO-0002572 | OA000-I07-S03-C006 | SSH5_pver | SVD5_I07 |  | x |  |  |
| I07S03C010POC | TARA_CO-0002576 | OA000-I07-S03-C010 | *P. effusa* | SVD1_I07 |  | x | A+B | x |
| I08S01C001POC | TARA_CO-0002596 | OA000-I08-S01-C001 | *P. verrucosa* | SVD3_I08 |  | x | B |  |
| I08S01C006POC | TARA_CO-0002601 | OA000-I08-S01-C006 | *P. meandrina* | SVD2_I08 |  | x | B |  |
| I08S01C010POC | TARA_CO-0002605 | OA000-I08-S01-C010 | *P. verrucosa* | SVD3_I08 |  | x |  |  |
| I08S02C001POC | TARA_CO-0002686 | OA000-I08-S02-C001 | *P. verrucosa* | SVD3_I08 |  | x |  |  |
| I08S02C006POC | TARA_CO-0002691 | OA000-I08-S02-C006 | *P. meandrina* | SVD2_I08 |  | x |  |  |
| I08S02C010POC | TARA_CO-0002695 | OA000-I08-S02-C010 | *P. meandrina* | SVD2_I08 |  | x | A |  |
| I08S03C001POC | TARA_OA-0002257 | OA000-I08-S03-C001 | *P. verrucosa* | SVD3_I08 |  | x | A |  |
| I08S03C006POC | TARA_OA-0002262 | OA000-I08-S03-C006 | *P. meandrina* | SVD2_I08 |  | x |  |  |
| I08S03C010POC | TARA_OA-0002266 | OA000-I08-S03-C010 | *P. verrucosa* | SVD3_I08 |  | x |  |  |
| I09S01C001POC | TARA_OA-0002323 | OA000-I09-S01-C001 | *P. verrucosa* | SVD3_I09 |  | x | B |  |
| I09S01C006POC | TARA_OA-0002328 | OA000-I09-S01-C006 | *P. meandrina* | SVD2_I09 |  | x | B |  |
| I09S01C010POC | TARA_OA-0002332 | OA000-I09-S01-C010 | *P. meandrina* | SVD2_I09 |  | x |  | x |
| I09S02C001POC | TARA_FH-0000951 | OA000-I09-S02-C001 | *P. verrucosa* | SVD3_I09 |  | x |  |  |
| I09S02C006POC | TARA_FH-0000956 | OA000-I09-S02-C006 | *P. verrucosa* | SVD3_I09 |  | x | A |  |
| I09S02C010POC | TARA_FH-0000960 | OA000-I09-S02-C010 | *P. verrucosa* | SVD3_I09 |  | x |  |  |
| I09S03C001POC | TARA_OA-0002451 | OA000-I09-S03-C001 | *P. meandrina* | SVD2_I09 |  | x | A |  |
| I09S03C006POC | TARA_OA-0002456 | OA000-I09-S03-C006 | *P. verrucosa* | SVD3_I09 |  | x |  | x |
| I09S03C010POC | TARA_OA-0002460 | OA000-I09-S03-C010 | SVD2 SVD3 | SVD2 SVD3_I09 |  | x | A+B |  |
| I10S01C001POC | TARA_CO-0004051 | OA000-I10-S01-C001 | *P. meandrina* | SVD2_I10 |  | x |  |  |
| I10S01C006POC | TARA_CO-0004036 | OA000-I10-S01-C006 | *P. verrucosa* | SVD3_I10 |  | x | B |  |
| I10S01C010POC | TARA_CO-0004040 | OA000-I10-S01-C010 | *P. meandrina* | SVD2_I10 |  | x | B |  |
| I10S02C001POC | TARA_CO-0004201 | OA000-I10-S02-C001 | *P. meandrina* | SVD2_I10 |  | x | A |  |
| I10S02C006POC | TARA_CO-0004206 | OA000-I10-S02-C006 | *P. meandrina* | SVD2_I10 |  | x |  |  |
| I10S02C010POC | TARA_CO-0004210 | OA000-I10-S02-C010 | *P. verrucosa* | SVD3_I10 |  | x | A | x |
| I10S03C001POC | TARA_CO-0004321 | OA000-I10-S03-C001 | *P. meandrina* | SVD2_I10 |  | x |  |  |
| I10S03C006POC | TARA_CO-0004326 | OA000-I10-S03-C006 | *P. meandrina* | SVD2_I10 |  | x |  |  |
| I10S03C010POC | TARA_CO-0004330 | OA000-I10-S03-C010 | *P. meandrina* | SVD2_I10 |  | x |  | x |
| I15S01C001POC | TARA_CO-0001811 | OA000-I15-S01-C001 | *P. meandrina* | SVD2_I15 |  | x | B |  |
| I15S01C006POC | TARA_CO-0001816 | OA000-I15-S01-C006 | *P. verrucosa* | SVD3_I15 |  | x | B |  |
| I15S01C010POC | TARA_CO-0001820 | OA000-I15-S01-C010 | *P. verrucosa* | SVD3_I15 |  | x |  |  |
| I15S02C001POC | TARA_CO-0001659 | OA000-I15-S02-C001 | *P. verrucosa* | SVD3_I15 |  | x | A |  |
| I15S02C006POC | TARA_CO-0001664 | OA000-I15-S02-C006 | *P. meandrina* | SVD2_I15 |  | x |  |  |
| I15S02C010POC | TARA_CO-0001668 | OA000-I15-S02-C010 | *P. verrucosa* | SVD3_I15 |  | x |  |  |
| I15S03C001POC | TARA_CO-0001599 | OA000-I15-S03-C001 | *P. verrucosa* | SVD3_I15 |  | x |  |  |
| I15S03C006POC | TARA_CO-0001604 | OA000-I15-S03-C006 | *P. meandrina* | SVD2_I15 |  | x | A |  |
| I15S03C010POC | TARA_CO-0001608 | OA000-I15-S03-C010 | *P. verrucosa* | SVD3_I15 |  | x |  |  |
| **sample name** | **TARA barcode** | **sampling-design_label** | **SSH,subcluster** | **introgression population** | **MLL (clone)** | **ramets kept** | **BFD*** |  |
| ***Porites*** |  |  |  |  |  |  |  |  |
| I01S01C011POR | TARA_CO-0000035 | OA000-I01-S01-C011 | SSH1,K1a | K1a_I01 |  |  | A |  |
| I01S01C012POR | TARA_CO-0000038 | OA000-I01-S01-C012 | SSH1,K1a | K1a_I01 |  |  |  |  |
| I01S01C016POR | TARA_CO-0000050 | OA000-I01-S01-C016 | SSH1,K1a | K1a_I01 |  |  |  |  |
| I01S02C011POR | TARA_CO-0000075 | OA000-I01-S02-C011 | SSH1,K1a | K1a_I01 |  |  |  |  |
| I01S02C012POR | TARA_CO-0000076 | OA000-I01-S02-C012 | SSH1,K1a | K1a_I01 |  |  |  |  |
| I01S02C016POR | TARA_CO-0000080 | OA000-I01-S02-C016 | SSH1,K1a | K1a_I01 |  |  |  |  |
| I01S03C011POR | TARA_CO-0000505 | OA000-I01-S03-C011 | SSH1,K1a | K1a_I01 |  |  |  |  |
| I01S03C012POR | TARA_CO-0000506 | OA000-I01-S03-C012 | SSH1,K1a | K1a_I01 |  |  |  |  |
| I01S03C016POR | TARA_CO-0000510 | OA000-I01-S03-C016 | SSH1,K1a | K1a_I01 |  |  | B |  |
| I02S01C011POR | TARA_CO-0000150 | OA000-I02-S01-C011 | SSH2,K2a | K2a_I02 |  |  | A |  |
| I02S01C012POR | TARA_CO-0000151 | OA000-I02-S01-C012 | SSH2,K2a | K2a_I02 |  |  |  |  |
| I02S01C016POR | TARA_CO-0000155 | OA000-I02-S01-C016 | SSH1,K1a | K1a_I02 |  |  | A |  |
| I02S02C011POR | TARA_CO-0000230 | OA000-I02-S02-C011 | SSH2,K2a | K2a_I02 |  |  |  |  |
| I02S02C012POR | TARA_CO-0000231 | OA000-I02-S02-C012 | SSH2,K2a | K2a_I02 |  |  | B |  |
| I02S02C016POR | TARA_CO-0000235 | OA000-I02-S02-C016 | SSH2,K2a | K2a_I02 |  |  |  |  |
| I02S03C011POR | TARA_CO-0000395 | OA000-I02-S03-C011 | SSH1,K1a | K1a_I02 |  |  | B |  |
| I02S03C012POR | TARA_CO-0000396 | OA000-I02-S03-C012 | SSH2,K2a | K2a_I02 |  |  | A |  |
| I02S03C016POR | TARA_CO-0000400 | OA000-I02-S03-C016 | SSH2,K2a | K2a_I02 |  |  | B |  |
| I03S01C001POR | TARA_CO-0000525 | OA000-I03-S01-C001 | SSH1,K1b | K1b_I03 |  |  | A |  |
| I03S01C006POR | TARA_CO-0000530 | OA000-I03-S01-C006 | SSH1,K1a | K1a_I03 |  |  | A+B |  |
| I03S01C010POR | TARA_CO-0000534 | OA000-I03-S01-C010 | SSH1,K1b | K1b_I03 |  |  | B |  |
| I03S01C021POR | TARA_CO-0000565 | OA000-I03-S01-C021 | SSH1,K1b | K1b_I03 |  |  |  |  |
| I03S01C022POR | TARA_CO-0000566 | OA000-I03-S01-C022 | SSH1,K1b | K1b_I03 |  |  |  |  |
| I03S01C024POR | TARA_CO-0000568 | OA000-I03-S01-C024 | SSH1,K1b | K1b_I03 |  |  |  |  |
| I03S01C025POR | TARA_CO-0000569 | OA000-I03-S01-C025 | SSH1,K1b | K1b_I03 |  |  |  |  |
| I03S01C026POR | TARA_CO-0000570 | OA000-I03-S01-C026 | SSH1,K1b | K1b_I03 |  |  |  |  |
| I03S01C027POR | TARA_CO-0000571 | OA000-I03-S01-C027 | SSH1,K1b | K1b_I03 |  |  |  |  |
| I03S01C028POR | TARA_CO-0000572 | OA000-I03-S01-C028 | SSH1,K1b | K1b_I03 |  |  |  |  |
| I03S01C029POR | TARA_CO-0000573 | OA000-I03-S01-C029 | SSH1,K1b | K1b_I03 |  |  | A |  |
| I03S01C030POR | TARA_CO-0000574 | OA000-I03-S01-C030 | SSH1,K1b | K1b_I03 |  |  | B |  |
| I04S01C011POR | TARA_CO-0000590 | OA000-I04-S01-C011 | SSH3,K3a | K3a_I04 |  |  | A |  |
| I04S01C012POR | TARA_CO-0000591 | OA000-I04-S01-C012 | SSH3,K3a | K3a_I04 |  |  | B |  |
| I04S01C016POR | TARA_CO-0000595 | OA000-I04-S01-C016 | SSH3,K3a | K3a_I04 |  |  |  |  |
| I04S02C011POR | TARA_CO-0000675 | OA000-I04-S02-C011 | SSH3,K3a | K3a_I04 |  |  |  |  |
| I04S02C012POR | TARA_CO-0000676 | OA000-I04-S02-C012 | SSH3,K3a | K3a_I04 |  |  |  |  |
| I04S02C016POR | TARA_CO-0000680 | OA000-I04-S02-C016 | SSH3,K3a | K3a_I04 |  |  |  |  |
| I04S03C011POR | TARA_CO-0000727 | OA000-I04-S03-C011 | SSH3,K3a | K3a_I04 |  |  |  |  |
| I04S03C012POR | TARA_CO-0000728 | OA000-I04-S03-C012 | SSH3,K3a | K3a_I04 |  |  |  |  |
| I04S03C016POR | TARA_CO-0000732 | OA000-I04-S03-C016 | SSH3,K3a | K3a_I04 |  |  |  |  |
| I04S04C011POR | TARA_CO-0000787 | OA000-I04-S04-C011 | SSH3,K3a | K3a_I04 |  |  | A |  |
| I04S04C012POR | TARA_CO-0000788 | OA000-I04-S04-C012 | SSH3,K3a | K3a_I04 |  |  |  |  |
| I04S04C016POR | TARA_CO-0000792 | OA000-I04-S04-C016 | SSH3,K3a | K3a_I04 |  |  | B |  |
| I05S01C011POR | TARA_CO-0000797 | OA000-I05-S01-C011 | SSH3,K3b | K3b_I05 |  |  | A |  |
| I05S01C012POR | TARA_CO-0000798 | OA000-I05-S01-C012 | SSH3,K3b | K3b_I05 |  |  | B |  |
| I05S01C016POR | TARA_CO-0000802 | OA000-I05-S01-C016 | SSH2,K2b | K2b_I05 |  |  | A |  |
| I05S02C011POR | TARA_CO-0000908 | OA000-I05-S02-C011 | SSH3,K3b | K3b_I05 |  |  |  |  |
| I05S02C012POR | TARA_CO-0000909 | OA000-I05-S02-C012 | SSH3,K3b | K3b_I05 |  |  |  |  |
| I05S02C016POR | TARA_CO-0000913 | OA000-I05-S02-C016 | SSH3,K3b | K3b_I05 |  |  |  |  |
| I05S03C011POR | TARA_CO-0000958 | OA000-I05-S03-C011 | SSH3,K3b | K3b_I05 |  |  | A |  |
| I05S03C012POR | TARA_CO-0000959 | OA000-I05-S03-C012 | SSH2,K2b | K2b_I05 |  |  | B |  |
| I05S03C016POR | TARA_CO-0000963 | OA000-I05-S03-C016 | SSH3,K3b | K3b_I05 |  |  | B |  |
| I06S01C011POR | TARA_CO-0002046 | OA000-I06-S01-C011 | SSH2,K2b | K2b_I06 |  |  | A |  |
| I06S01C012POR | TARA_CO-0002047 | OA000-I06-S01-C012 | SSH3,K3c | K3c_I06 |  |  | A |  |
| I06S01C016POR | TARA_CO-0002051 | OA000-I06-S01-C016 | SSH3,K3c | K3c_I06 |  |  |  |  |
| I06S02C011POR | TARA_CO-0002136 | OA000-I06-S02-C011 | SSH3,K3c | K3c_I06 |  |  |  |  |
| I06S02C012POR | TARA_CO-0002137 | OA000-I06-S02-C012 | SSH3,K3c | K3c_I06 |  |  |  |  |
| I06S02C016POR | TARA_CO-0002141 | OA000-I06-S02-C016 | SSH3,K3c | K3c_I06 |  |  |  |  |
| I06S03C011POR | TARA_CO-0002276 | OA000-I06-S03-C011 | SSH2,K2b | K2b_I06 |  |  | B |  |
| I06S03C012POR | TARA_CO-0002277 | OA000-I06-S03-C012 | SSH3,K3c | K3c_I06 |  |  | B |  |
| I06S03C016POR | TARA_CO-0002281 | OA000-I06-S03-C016 | SSH3,K3c | K3c_I06 |  |  |  |  |
| I07S01C011POR | TARA_CO-0002366 | OA000-I07-S01-C011 | SSH2,K2b | K2b_I07 |  |  | B |  |
| I07S01C012POR | TARA_CO-0002367 | OA000-I07-S01-C012 | SSH2,K2b | K2b_I07 |  |  |  |  |
| I07S01C016POR | TARA_CO-0002371 | OA000-I07-S01-C016 | SSH3,K3c | K3c_I07 |  |  | B |  |
| I07S01C017POR | TARA_CO-0002372 | OA000-I07-S01-C017 | SSH2,K2b | K2b_I07 |  |  |  |  |
| I07S02C011POR | TARA_CO-0002446 | OA000-I07-S02-C011 | SSH3,K3c | K3c_I07 |  |  |  |  |
| I07S02C012POR | TARA_CO-0002447 | OA000-I07-S02-C012 | SSH3,K3c | K3c_I07 |  |  |  |  |
| I07S02C013POR | TARA_CO-0002448 | OA000-I07-S02-C013 | SSH3,K3c | K3c_I07 |  |  |  |  |
| I07S02C014POR | TARA_CO-0002449 | OA000-I07-S02-C014 | SSH3,K3c | K3c_I07 |  |  | A |  |
| I07S02C016POR | TARA_CO-0002451 | OA000-I07-S02-C016 | SSH3,K3c | K3c_I07 |  |  |  |  |
| I07S03C011POR | TARA_CO-0002577 | OA000-I07-S03-C011 | SSH2,K2b | K2b_I07 |  |  |  |  |
| I07S03C012POR | TARA_CO-0002578 | OA000-I07-S03-C012 | SSH2,K2b | K2b_I07 |  |  |  |  |
| I07S03C016POR | TARA_CO-0002582 | OA000-I07-S03-C016 | SSH2,K2b | K2b_I07 |  |  |  |  |
| I07S03C018POR | TARA_CO-0002584 | OA000-I07-S03-C018 | SSH2,K2b | K2b_I07 |  |  | A |  |
| I08S01C011POR | TARA_CO-0002606 | OA000-I08-S01-C011 | SSH2,K2b | K2b_I08 |  |  | A |  |
| I08S01C012POR | TARA_CO-0002607 | OA000-I08-S01-C012 | SSH2,K2b | K2b_I08 |  |  |  |  |
| I08S01C016POR | TARA_CO-0002611 | OA000-I08-S01-C016 | SSH3,K3c | K3c_I08 |  |  | A |  |
| I08S02C011POR | TARA_CO-0002696 | OA000-I08-S02-C011 | SSH3,K3c | K3c_I08 |  |  |  |  |
| I08S02C012POR | TARA_CO-0002697 | OA000-I08-S02-C012 | SSH2,K2b | K2b_I08 |  |  | B |  |
| I08S02C016POR | TARA_CO-0002701 | OA000-I08-S02-C016 | SSH3,K3c | K3c_I08 |  |  |  |  |
| I08S03C011POR | TARA_OA-0002267 | OA000-I08-S03-C011 | SSH3,K3c | K3c_I08 |  |  |  |  |
| I08S03C012POR | TARA_OA-0002268 | OA000-I08-S03-C012 | SSH3,K3c | K3c_I08 |  |  |  |  |
| I08S03C016POR | TARA_OA-0002272 | OA000-I08-S03-C016 | SSH3,K3c | K3c_I08 |  |  | B |  |
| I09S01C011POR | TARA_OA-0002333 | OA000-I09-S01-C011 | SSH3,K3d | K3d_I09 |  |  | A |  |
| I09S01C012POR | TARA_OA-0002334 | OA000-I09-S01-C012 | SSH2,K2b | K2b_I09 |  |  | A |  |
| I09S01C016POR | TARA_OA-0002338 | OA000-I09-S01-C016 | SSH3,K3d | K3d_I09 |  |  |  |  |
| I09S02C011POR | TARA_FH-0000961 | OA000-I09-S02-C011 | SSH3,K3d | K3d_I09 |  |  |  |  |
| I09S02C012POR | TARA_FH-0000962 | OA000-I09-S02-C012 | SSH2,K2b | K2b_I09 |  |  | B |  |
| I09S02C016POR | TARA_FH-0000966 | OA000-I09-S02-C016 | SSH3,K3d | K3d_I09 |  |  |  |  |
| I09S03C011POR | TARA_OA-0002461 | OA000-I09-S03-C011 | SSH3,K3d | K3d_I09 |  |  |  |  |
| I09S03C012POR | TARA_OA-0002462 | OA000-I09-S03-C012 | SSH3,K3d | K3d_I09 |  |  |  |  |
| I09S03C016POR | TARA_OA-0002466 | OA000-I09-S03-C016 | SSH3,K3d | K3d_I09 |  |  | B |  |
| I10S01C011POR | TARA_CO-0004041 | OA000-I10-S01-C011 | SSH2,K2c | K2c_I10 |  |  | A |  |
| I10S01C012POR | TARA_CO-0004042 | OA000-I10-S01-C012 | SSH2,K2c | K2c_I10 |  |  |  |  |
| I10S01C016POR | TARA_CO-0004046 | OA000-I10-S01-C016 | SSH2,K2c | K2c_I10 |  |  |  |  |
| I10S02C011POR | TARA_CO-0004211 | OA000-I10-S02-C011 | SSH2,K2c | K2c_I10 |  |  |  |  |
| I10S02C012POR | TARA_CO-0004212 | OA000-I10-S02-C012 | SSH2,K2c | K2c_I10 |  |  |  |  |
| I10S02C016POR | TARA_CO-0004216 | OA000-I10-S02-C016 | SSH2,K2c | K2c_I10 |  |  |  |  |
| I10S03C011POR | TARA_CO-0004331 | OA000-I10-S03-C011 | SSH2,K2c | K2c_I10 |  |  |  |  |
| I10S03C012POR | TARA_CO-0004332 | OA000-I10-S03-C012 | SSH2,K2c | K2c_I10 |  |  |  |  |
| I10S03C016POR | TARA_CO-0004336 | OA000-I10-S03-C016 | SSH2,K2c | K2c_I10 |  |  | B |  |
| I15S01C011POR | TARA_CO-0001821 | OA000-I15-S01-C011 | SSH3,K3d | K3d_I15 |  |  | A |  |
| I15S01C012POR | TARA_CO-0001822 | OA000-I15-S01-C012 | SSH3,K3d | K3d_I15 |  |  |  |  |
| I15S01C016POR | TARA_CO-0001826 | OA000-I15-S01-C016 | SSH2,K2c | K2c_I15 |  |  | A |  |
| I15S02C011POR | TARA_CO-0001669 | OA000-I15-S02-C011 | K2/K3,K2/K3 | K2/K3_I15 |  |  | A+B |  |
| I15S02C012POR | TARA_CO-0001670 | OA000-I15-S02-C012 | SSH2,K2c | K2c_I15 |  |  |  |  |
| I15S02C016POR | TARA_CO-0001674 | OA000-I15-S02-C016 | SSH2,K2c | K2c_I15 |  |  | B |  |
| I15S03C011POR | TARA_CO-0001609 | OA000-I15-S03-C011 | SSH3,K3d | K3d_I15 |  |  |  |  |
| I15S03C012POR | TARA_CO-0001610 | OA000-I15-S03-C012 | SSH3,K3d | K3d_I15 |  |  | B |  |
| I15S03C016POR | TARA_CO-0001614 | OA000-I15-S03-C016 | SSH2,K2c | K2c_I15 |  |  | A+B |  |
| **sample name** | **TARA barcode** | **sampling-design_label** | **sNMF,SVDquartet** | **introgression population** | **MLL (clone)** | **ramets kept** |  |  |
| ***Millepora*** |  |  |  |  |  |  |  |  |
| I02S01C021MIL | TARA_CO-0000200 | OA000-I02-S01-C021 | K4,SVD1 | I02 | 1_54 | x |  |  |
| I02S01C022MIL | TARA_CO-0000201 | OA000-I02-S01-C022 | K4,SVD1 | I02 | 1_54 | x |  |  |
| I02S01C031MIL | TARA_CO-0000302 | OA000-I02-S01-C031 | K4,SVD1 | I02 | 1_54 | x |  |  |
| I06S01C021MIL | TARA_CO-0002036 | OA000-I06-S01-C021 | K2,SVD6 | I06 |  | x |  |  |
| I06S01C022MIL | TARA_CO-0002037 | OA000-I06-S01-C022 | K2,SVD6 | I06 |  | x |  |  |
| I06S01C028MIL | TARA_CO-0002043 | OA000-I06-S01-C028 | K2,SVD6 | I06 |  | x |  |  |
| I06S02C021MIL | TARA_CO-0002176 | OA000-I06-S02-C021 | K2,SVD6 | I06 | 1_46 |  |  |  |
| I06S02C022MIL | TARA_CO-0002177 | OA000-I06-S02-C022 | K2,SVD6 | I06 | 1_46 | x |  |  |
| I06S02C026MIL | TARA_CO-0002181 | OA000-I06-S02-C026 | K2,SVD6 | I06 | 1_46 |  |  |  |
| I06S03C021MIL | TARA_CO-0002266 | OA000-I06-S03-C021 | K2,SVD6 | I06 | 1_21 | x |  |  |
| I06S03C022MIL | TARA_CO-0002267 | OA000-I06-S03-C022 | K2,SVD6 | I06 | 1_21 |  |  |  |
| I06S03C026MIL | TARA_CO-0002271 | OA000-I06-S03-C026 | K2,SVD6 | I06 |  | x |  |  |
| I07S01C021MIL | TARA_CO-0002376 | OA000-I07-S01-C021 | K3,SVD4 | I07 |  | x |  |  |
| I07S01C022MIL | TARA_CO-0002377 | OA000-I07-S01-C022 | K3,SVD4 | I07 |  | x |  |  |
| I07S01C026MIL | TARA_CO-0002381 | OA000-I07-S01-C026 | K3,SVD4 | I07 |  | x |  |  |
| I07S02C021MIL | TARA_CO-0002426 | OA000-I07-S02-C021 | K3,SVD4 | I07 |  | x |  |  |
| I07S02C022MIL | TARA_CO-0002427 | OA000-I07-S02-C022 | K3,SVD4 | I07 |  | x |  |  |
| I07S02C026MIL | TARA_CO-0002431 | OA000-I07-S02-C026 | K3,SVD4 | I07 |  | x |  |  |
| I07S03C021MIL | TARA_CO-0002587 | OA000-I07-S03-C021 | K3,SVD4 | I07 |  | x |  |  |
| I07S03C022MIL | TARA_CO-0002588 | OA000-I07-S03-C022 | K3,SVD4 | I07 |  | x |  |  |
| I07S03C026MIL | TARA_CO-0002592 | OA000-I07-S03-C026 | K3,SVD4 | I07 |  | x |  |  |
| I08S01C021MIL | TARA_CO-0002617 | OA000-I08-S01-C021 | K2,SVD6 | I08 |  | x |  |  |
| I08S01C022MIL | TARA_CO-0002618 | OA000-I08-S01-C022 | K2,SVD6 | I08 |  | x |  |  |
| I08S01C026MIL | TARA_CO-0002622 | OA000-I08-S01-C026 | K2,SVD6 | I08 |  | x |  |  |
| I08S02C021MIL | TARA_CO-0002706 | OA000-I08-S02-C021 | K2,SVD6 | I08 |  | x |  |  |
| I08S02C022MIL | TARA_CO-0002707 | OA000-I08-S02-C022 | K2,SVD6 | I08 |  | x |  |  |
| I08S02C026MIL | TARA_CO-0002711 | OA000-I08-S02-C026 | K2,SVD6 | I08 |  | x |  |  |
| I08S03C021MIL | TARA_OA-0002277 | OA000-I08-S03-C021 | K2,SVD6 | I08 |  | x |  |  |
| I08S03C022MIL | TARA_OA-0002278 | OA000-I08-S03-C022 | K2,SVD6 | I08 |  | x |  |  |
| I08S03C026MIL | TARA_OA-0002282 | OA000-I08-S03-C026 | K2,SVD6 | I08 |  | x |  |  |
| I09S01C021MIL | TARA_OA-0002343 | OA000-I09-S01-C021 | K1/K2,SVD5 | I09 |  | x |  |  |
| I09S01C022MIL | TARA_OA-0002344 | OA000-I09-S01-C022 | K1/K2,SVD5 | I09 | 1_33 | x |  |  |
| I09S01C026MIL | TARA_OA-0002348 | OA000-I09-S01-C026 | K1/K2,SVD5 | I09 | 1_33 |  |  |  |
| I09S02C021MIL | TARA_FH-0000971 | OA000-I09-S02-C021 | K1/K2,SVD5 | I09 |  | x |  |  |
| I09S02C022MIL | TARA_FH-0000972 | OA000-I09-S02-C022 | K1/K2,SVD5 | I09 |  | x |  |  |
| I09S02C026MIL | TARA_FH-0000976 | OA000-I09-S02-C026 | K1/K2,SVD5 | I09 |  | x |  |  |
| I09S03C021MIL | TARA_OA-0002471 | OA000-I09-S03-C021 | K1/K2,SVD5 | I09 |  | x |  |  |
| I09S03C022MIL | TARA_OA-0002472 | OA000-I09-S03-C022 | K1/K2,SVD5 | I09 |  | x |  |  |
| I10S01C021MIL | TARA_CO-0004031 | OA000-I10-S01-C021 | K1/K3/K2,SVD3 | I10 |  | x |  |  |
| I10S01C022MIL | TARA_CO-0004032 | OA000-I10-S01-C022 | K1/K3/K2,SVD3 | I10 |  | x |  |  |
| I10S01C026MIL | TARA_CO-0004056 | OA000-I10-S01-C026 | K1/K3/K2,SVD3 | I10 |  | x |  |  |
| I10S02C021MIL | TARA_CO-0004221 | OA000-I10-S02-C021 | K1/K3/K2,SVD3 | I10 |  | x |  |  |
| I10S02C022MIL | TARA_CO-0004222 | OA000-I10-S02-C022 | K1/K3/K2,SVD3 | I10 |  | x |  |  |
| I10S02C026MIL | TARA_CO-0004226 | OA000-I10-S02-C026 | K1/K3/K2,SVD3 | I10 |  | x |  |  |
| I10S03C021MIL | TARA_CO-0004341 | OA000-I10-S03-C021 | K1/K3/K2,SVD3 | I10 |  | x |  |  |
| I10S03C022MIL | TARA_CO-0004342 | OA000-I10-S03-C022 | K1/K3/K2,SVD3 | I10 |  | x |  |  |
| I10S03C026MIL | TARA_CO-0004346 | OA000-I10-S03-C026 | K1/K3/K2,SVD3 | I10 |  | x |  |  |
| I15S01C021MIL | TARA_CO-0001831 | OA000-I15-S01-C021 | K1,SVD2 | I15 |  | x |  |  |
| I15S01C022MIL | TARA_CO-0001832 | OA000-I15-S01-C022 | K1,SVD2 | I15 |  | x |  |  |
| I15S01C026MIL | TARA_CO-0001836 | OA000-I15-S01-C026 | K1,SVD2 | I15 |  | x |  |  |
| I15S02C021MIL | TARA_CO-0001679 | OA000-I15-S02-C021 | K1,SVD2 | I15 | 1_40 |  |  |  |
| I15S02C022MIL | TARA_CO-0001682 | OA000-I15-S02-C022 | K1,SVD2 | I15 | 1_40 | x |  |  |
| I15S02C026MIL | TARA_CO-0001685 | OA000-I15-S02-C026 | K1,SVD2 | I15 |  | x |  |  |
| I15S03C021MIL | TARA_CO-0001619 | OA000-I15-S03-C021 | K1,SVD2 | I15 |  | x |  |  |
| I15S03C022MIL | TARA_CO-0001620 | OA000-I15-S03-C022 | K1,SVD2 | I15 |  | x |  |  |
| I15S03C026MIL | TARA_CO-0001624 | OA000-I15-S03-C026 | K1,SVD2 | I15 |  | x |  |  |
