## Supplementary_Table_4 for "Disparate patterns of genetic divergence in three widespread corals across a pan-Pacific environmental gradient highlights species-specific adaptation trajectories"

**Supplementary Table 4.** **Mantel tests of genetic distances against geographic and historical temperature distances**. Tests are given for each of the genera and for each of the lineages within the genera. Tests for correlation between geographic and historical temperature distances for each of the genera are also given. Significant (p<0.05) results are underlined.

|  | **Distance** | **Mantel’s r** | **p value** |
| --- | --- | --- | --- |
| **Genera** |  |  |  |
| *Pocillopora* | geographic | -0.019 | 0.517 |
|  | environmental | -0.044 | 0.568 |
| *Porites* | geographic | 0.397 | 0.005 |
|  | environmental | 0.361 | 0.020 |
| *Millepora* | geographic | 0.722 | 0.053 |
|  | environmental | 0.930 | 0.019 |
| **Lineage** |  |  |  |
| *Pocillopora* |  |  |  |
| SVD1 | geographic | -0.376 | 0.667 |
|  | environmental | -0.577 | 0.833 |
| SVD2 | geographic | 0.514 | 0.167 |
|  | environmental | -0.229 | 0.683 |
| SVD3 | geographic | 0.772 | 0.083 |
|  | environmental | -0.661 | 1.000 |
| SVD4 | geographic | 0.722 | 0.083 |
|  | environmental | -0.103 | 0.650 |
| SVD5 | geographic | 0.677 | 0.333 |
|  | environmental | 0.816 | 0.333 |
| *Porites* |  |  |  |
| K1 | geographic | 0.982 | 0.167 |
|  | environmental | -0.924 | 1.000 |
| K2 | geographic | 0.585 | 0.036 |
|  | environmental | 0.641 | 0.010 |
| K3 | geographic | 0.591 | 0.008 |
|  | environmental | 0.372 | 0.052 |
